# Seasonal Metabolic Dynamics of Microeukaryotic Plankton: A Year-long Metatranscriptomic Study in a Temperate Sea

**DOI:** 10.1101/2024.01.17.576024

**Authors:** Michiel Perneel, Rune Lagaisse, Jonas Mortelmans, Steven Maere, Pascal I. Hablützel

## Abstract

Seasonal fluctuations profoundly affect marine microeukaryotic plankton composition and metabolism, but accurately tracking these changes has been a longstanding challenge. In this study, we present a year-long metatranscriptomic dataset from the Southern Bight of the North Sea, shedding light on the seasonal dynamics in temperate plankton ecosystems. We observe distinct shifts in active plankton species and their metabolic processes in response to seasonal changes. We characterised the metabolic signatures of different seasonal phases in detail, thereby revealing the metabolic versatility of dinoflagellates, the heterotrophic dietary strategy of *Phaeocystis* during its late-stage blooms, and diatoms being most abundant and metabolically active in autumn. Our data illuminates the varied contributions of microeukaryotic taxa to biomass production and nutrient cycling at different times of the year and allows to delineate their ecological niches. These findings underscore the use of metatranscriptomics for continuous marine ecosystem monitoring to enhance our ecological understanding of the ocean’s eukaryotic microbiome.

## Introduction

Marine microeukaryotic plankton play a pivotal role in primary production, global oceanic biogeochemical cycles, ecosystem stability, and climate regulation^1–4^. They exhibit pronounced spatiotemporal dynamics with recurring seasonal phenological patterns^5^. Such phenological rhythms, like the yearly blooms of both phototrophic microalgae and heterotrophic grazers, are particularly sensitive to disruption by anthropogenically altered environmental drivers^6–9^. In the face of current climate change, understanding the patterns and drivers of seasonal microeukaryote ecosystem dynamics, i.e. cyclic variations in species composition, biomass, and metabolic activity, is paramount for deciphering marine ecosystem functionality and resilience^10^.

Recent advances and maturation of methodological approaches in the marine sciences, including plankton abundance time series^11,12^, large-scale -omics surveys^13–16^, and expansion of genetic reference databases^17–20^, have been propelling a deeper understanding of marine ecosystems. However, many studies only track natural microeukaryotic plankton assemblages over limited periods of time^21^, or only at a few dates of the seasonal cycle^22^, whereas complete seasonal trajectories are needed to fully understand the temporal dynamics of plankton assemblages and the associated metabolic functions. Metatranscriptomic profiling provides an underused opportunity for a more comprehensive characterization of coastal microeukaryote plankton ecosystems^23–26^. The systematic capture of microeukaryotic plankton gene expression profiles over time from fixed locations generates ecosystem snapshots that facilitate the reconstruction of the turnover and relationships of active species and their metabolic responses to environmental fluctuations.

In this study, we aim to capture the phenological patterns and metabolic functions that follow seasonal turnover in temperate micro-eukaryotic plankton assemblages of the coastal Southern Bight of the North Sea. This shallow marginal sea is characterised by a complex subtidal sand bank system and high nutrient input from the Rhine-Meuse-Scheldt estuary^27^. We generated a year-long time series of metatranscriptomic data, which elucidates the unique functional properties of various ecosystem states and reveals typical metabolic traits of dominant plankton groups. Additionally, we investigate key features of the Southern Bight plankton assemblages such as the relationship between biodiversity and functional richness, the spatial variation resulting from the mixing of oceanic and estuarine waters, and seasonal shifts in feeding modes of certain microeukaryote species.

## Results

### The Belgian North Sea

Six locations in the Belgian North Sea were sampled monthly to construct time series of oceanographic, meteorological, nutrient, pigment, biotic and metatranscriptomic data (see Methods, Supplementary Table 1, and Fig. 1). Water temperature in the southern North Sea measured between 2.17 and 22.54 °C in winter and in summer, respectively. Nutrient concentrations rose over winter, reaching the highest values in February, and were depleted by April (Supplementary Dataset 1). Suspended particulate matter (SPM) and salinity also varied, with fresher and more turbid water in winter, and more saline and clearer water in summer/fall. Spatially, levels of SPM, nitrate, phosphate, and silicate differed across locations, with stations 130 and 700 often showing the highest concentrations (Fig. 1, Supplementary Dataset 1). Spatial differences in nutrient load and salinity in the study area are structured according to proximity to the Scheldt estuary in the east, with its freshwater discharge decreasing salinity and increasing nutrient levels in the Belgian coastal waters, while the inflowing North Atlantic water through the English channel from the west buffers temperature variations^28^.

**Figure 1.**
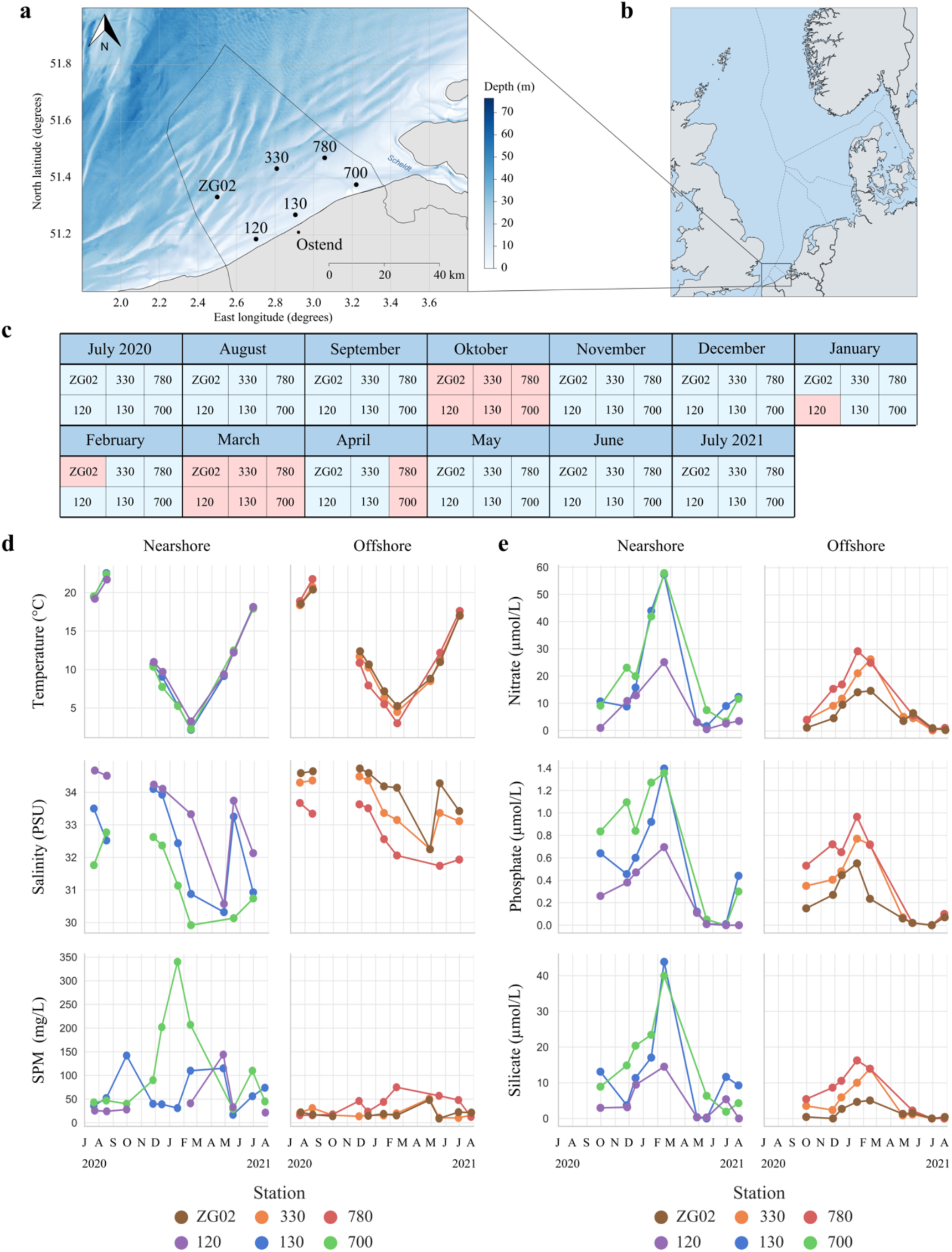
Sampling stations and their temporal and oceanographic context. **a)** Geographic location of the six sampling locations in the Belgian North Sea*^70^*. **b)** Location of the sampling area in the wider North Sea region (map by NordNordWest/Wikipedia, distributed under a CC-BY-SA-3.0-DE license). **c)** From July 2020 to July 2021, metatranscriptomic data was generated from these 6 stations. Gaps (in red) are due to cancelled campaigns because of stormy weather conditions or COVID-19 measures. For October 2020 and March 2021, weather conditions were too harsh to visit any station. **d)** Seasonal changes in temperature, salinity and suspended particulate matter (SPM) for near- and offshore stations. **e)** Seasonal changes in nitrate, phosphate, and silicate nutrient levels for near- and offshore stations.

### Phenology of taxonomic groups

Our metatranscriptomic analysis yielded over 1.049 billion raw reads and 7 million unique transcripts, functional or shallow taxonomic annotation information could be obtained for 79% of 3.7 million predicted proteins (see Fig. S1 and details in supplementary information). Diatoms were present year-round (Fig. 2), but different diatom assemblages succeeded each other (Fig. S2 & S3). *Phaeocystis* bloomed in April and was detected in 3 of the 4 stations visited that month (Fig. 3). From May to July, the microeukaryote ecosystem was characterised by high relative abundances of dinoflagellate transcripts, with *Noctiluca* being the most abundant dinoflagellate genus during those months. In fall and winter, arthropods were relatively more abundant. Some taxonomic groups only occurred in certain months, e.g. high relative abundances of the ctenophore *Mnemiopsis leidyi* were found in August. Similarly, most spirotrichs were found in August and September. Microplanktonic biomass, determined as cell densities per litre of seawater using FlowCam automated microscopy, peaked in July (Fig. 2c).

**Figure 2.**
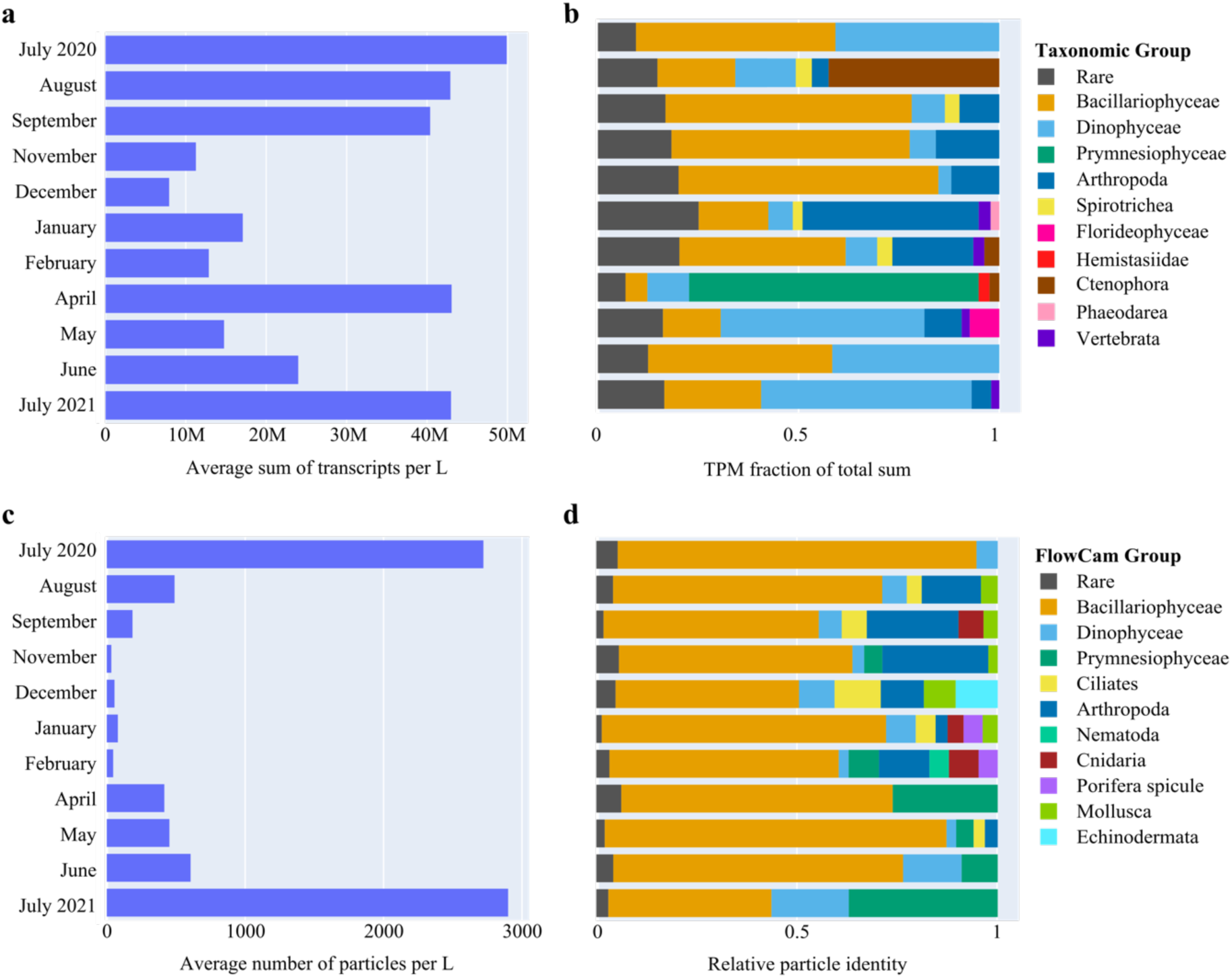
Monthly biomass fluctuations and turnover in taxonomic composition. **a)** Sum of transcripts per liter of sea surface water, averaged across samples in a month. **b)** Monthly relative abundances of high-level taxonomic groups annotated using EukProt (>60% sequence identity with reference), averaged over sampling stations. Relative abundance was calculated as the total sum of transcripts per million (TPM) of that group for a given sampling month, divided by the total TPM sum over all groups for that month (excluding unannotated transcripts). When relative abundance of a group was <2 %, they were labelled as ‘rare’. **c)** Average number of particles per liter of seawater, calculated using FlowCam automated image analysis. **d)** Relative cell densities of taxonomic groups per month, averaged over stations, as observed through FlowCam automated image classification. When relative abundance of a group was <2 %, they were labelled as ‘rare’.

**Figure 3.**
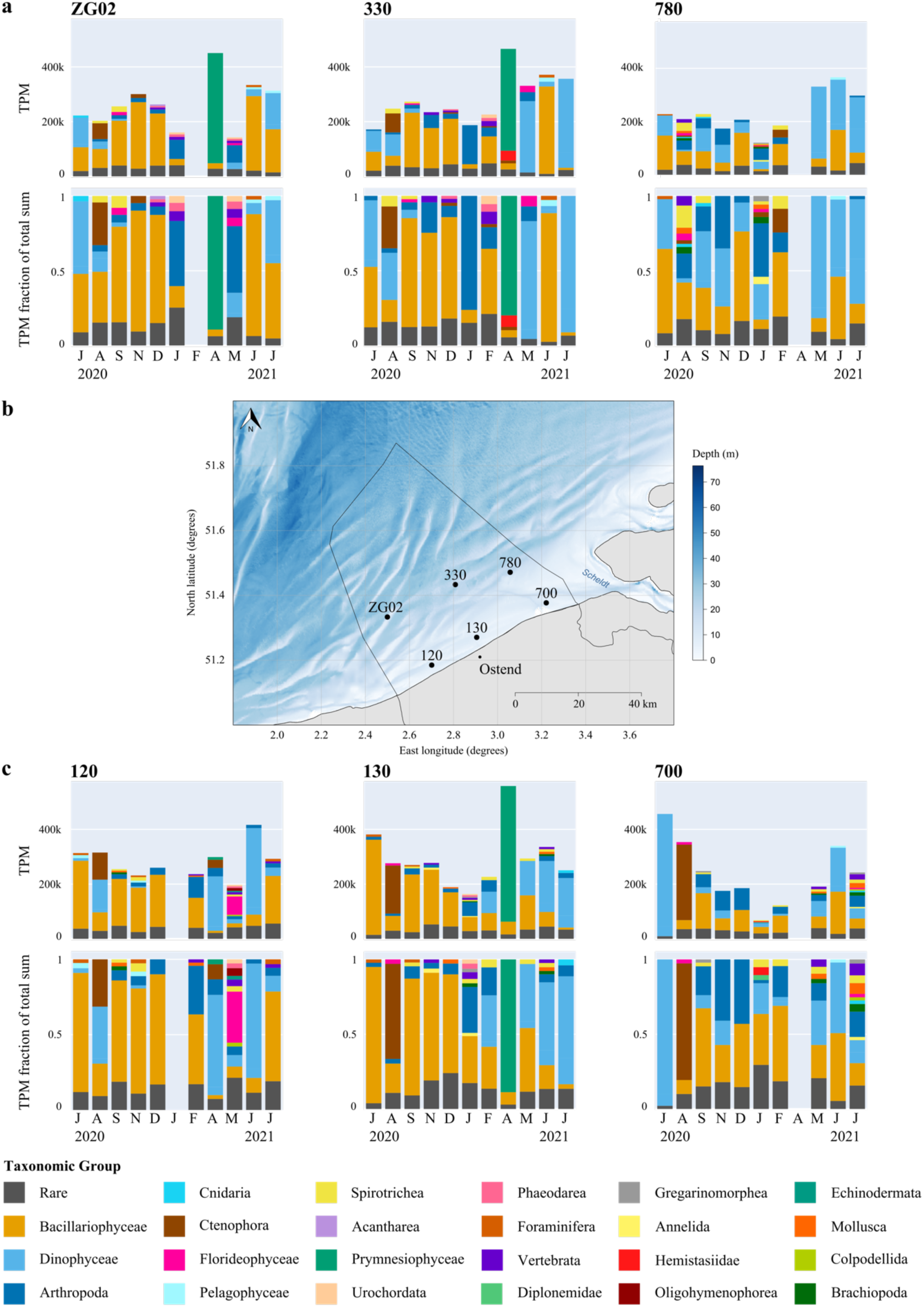
Monthly turnover in transcript abundance and relative taxonomic composition at each sampling station. **a)** Monthly absolute transcript abundance (top panels) and relative transcript abundance (bottom panels) for taxonomic groups annotated using EukProt, per offshore sampling station. Transcript abundance represents the sum of transcripts per million (TPM) belonging to a taxonomic group. Relative abundance was calculated as the sum of TPM for that group for a given sample, divided by the total TPM of all groups found in that sample. When relative abundance of a group was <2 %, they were labelled as ‘rare’. **b)** Spatial location of the 6 sampling stations in the Belgian North Sea. **c)** Monthly absolute transcript abundance (top panels) and relative transcript abundance (bottom panels) for taxonomic groups annotated using EukProt, per nearshore sampling station. Absolute and relative transcript abundances were calculated as for the offshore stations.

The spatial variability in taxonomic composition and transcript abundance across the six stations is substantial (Fig. 3). For example, we detected *Hemistasia*, a flagellate predator of *Phaeocystis* and other protists^29^, in only one of the three stations that had a *Phaeocystis* bloom. More generally, diatoms exhibited a lower relative expression at stations 780 and 700. The persistent spatial differences likely reflect the presence of a mixing zone where inflowing North Atlantic waters meet outflowing Rhine-Meuse-Scheldt waters, which creates a front zone that influences species dynamics and occurrences.

### Exploring Functional Diversity

Absolute transcript abundance was found to be highest when FlowCam-derived cell abundance levels were at their peak (Fig. 2, details in supplementary information). However, in April, there was a discrepancy between plankton biomass and transcript abundance, likely attributable to the challenge of detecting blooming *Phaeocystis* colonies using FlowCam imaging.

Functional richness, quantified as the number of unique enzymatic functions (KEGG KO identifiers) present in a sample, did not differ between months (Fig. S4; ANOVA, F(10,51) = 1.062, with degrees of freedom 10 and 51 representing #groups -1 and #samples - #groups, p = 0.40). Similarly, no significant differences were observed between stations (ANOVA, F(5,56) = 0.594, p = 0.704), or when considering the number of unique PFAM protein families instead (Fig. S5; ANOVA_months_, F(10,51) = 0.896, p = 0.544; ANOVA_stations_, F(5,56) = 0.731, p = 0.604). Principal component analysis of KEGG KO expression profiles however revealed temporal and spatial variation in the expression of metabolic functions in the microeukaryotic ecosystem across sampling months and stations (Fig. S4a). The functional dissimilarity between seasons coincides with distinct nutrient and temperature regimes, while the differences within summer months correlate with differences in suspended particulate matter load (Fig. S4 & Fig. 1a).

Active species richness averaged across stations was higher from September to February and lower from May to July (Fig. S6; ANOVA F(10,51) = 9.872, p = 6.185 * 10^-9^). No strong correlation was found between functional richness and the number of active species (Pearson’s correlation r = 0.18, df = 60, p = 0.151). A negative correlation was found between log transformed FlowCam cell abundances and the number of active species (r = -0.43, df = 60, p = 0.0008). Likewise, there is a weak negative correlation between functional richness and log transformed FlowCam cell abundances (r = -0.28, df = 55, p = 0.036). These negative correlations likely arise from non-homogeneous distribution of species in high biomass situations, which impacts the detection of rare species with the given sequencing depth^30^ (Fig. S6).

In conclusion, from September to February, there is an increase in the number of metabolically active species. This increase, however, does not result in increased functional richness.

### Functional seasonal dynamics of micro-eukaryotic plankton

To get a better understanding of how metabolic functions vary and covary in the Southern North Sea micro-eukaryotic plankton assemblages and how this is related to the presence of particular taxa, we applied weighted gene co-expression network analysis^31^ to the KEGG KO expression profiles across samples. 10 distinct modules of co-expressed functional identifiers were identified (Fig. 4). Analysis of module eigengene expression revealed fall and winter clusters (M3, M8, M9, M10) that are correlated with the presence of copepods and/or certain diatom genera, and spring/summer clusters that represent different ecosystem states: a *Phaeocystis* bloom in April (M1), a dinoflagellate dominance from May to August (M4-M6), diatom assemblages in June and July (M2), and the occurrence of *Mnemiopsis* in August (M7) (Fig. 4). For each module, we then determined characteristic KEGG pathways that were strongly represented in the module (see Methods) and quantified the contribution of the genus whose absolute abundance was most strongly correlated to the module’s eigengene expression to the overall expression of these pathways.

**Figure 4.**
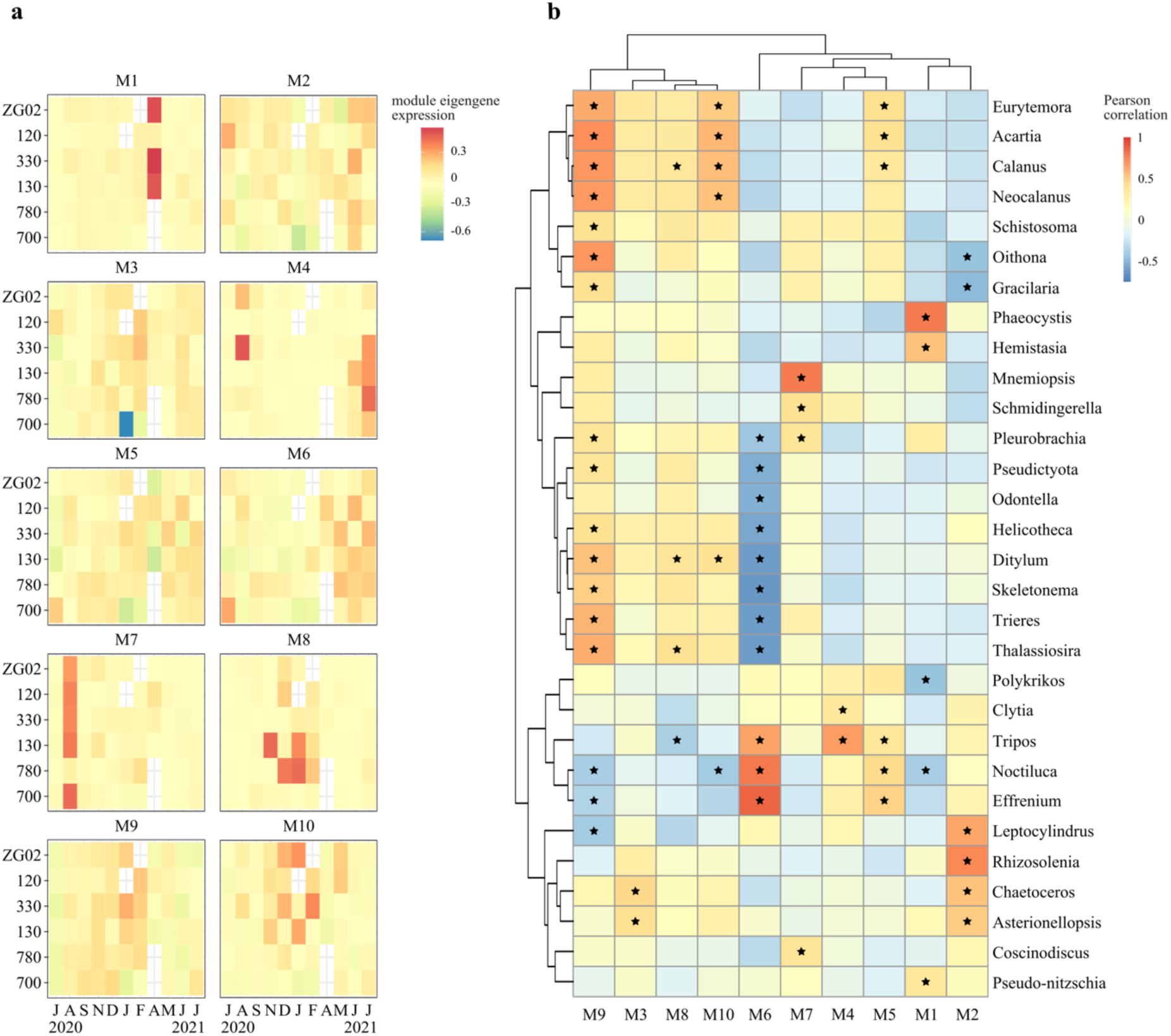
Module eigengene expression and correlation with taxonomic groups. **a)** Heatmap of module eigengenes per station per month. Eigengenes represent the first principal component of a module. Modules were calculated using weighted gene correlation network analysis (WGCNA) on transcript per million (TPM) expression values of KEGG KO identifiers. **b)** Pearson correlation of module eigengene expression per sample with log-transformed absolute genus abundances of EukProt-annotated transcripts (TPM sums per sample). Black stars indicate correlations with p-values < 0.05, corrected for multiple testing using the Benjamini & Hochberg correction method.

Module M1 eigengene expression predominantly correlates with increased absolute abundance of the prymnesiophyte genus *Phaeocystis* (Fig. 4b). KOs involved in cell cycle control and lysine and brassinosteroids biosynthesis displayed significantly higher ranked correlations with M1 eigengene expression than KOs associated with other pathways (Supplementary Dataset 2). For lysine biosynthesis and cell cycle control, most taxonomically annotated transcripts that were expressed during the April *Phaeocystis* bloom were indeed annotated to *Phaeocystis* (Fig. S7, Supplementary Dataset 2 & 3). None of the transcripts associated to brassinosteroids biosynthesis were annotated as *Phaeocystis*. This indicates that either the *Phaeocystis* reference transcriptome did not include genuine *Phaeocystis* genes involved in the biosynthesis of brassinosteroids, or that another unidentified organism produces brassinosteroids during a *Phaeocystis* bloom. Brassinosteroids have been associated with growth, development and protection against abiotic stress in plants, and could have similar functions in *Phaeocystis*^32^.

The module M2 eigengene profile exhibits a correlation with a spring/summer diatom assemblage, mainly with the genera *Rhizosolenia*, *Leptocylindrus*, *Chaetoceros*, and *Asterionellopsis* (Fig. 4). Functionally, this module contains KOs involved in the biosynthesis of amino acids and co-factors, biotin, and vitamin B6 metabolism (Fig. S7, Supplementary Dataset 2). Additionally, the identification of pathways related to porphyrin metabolism may imply regulation of pigments in response to light and nutrient availability. Taxonomic attribution of these pathways’ expression indicates a diatom contribution in fall, but not in spring (Fig. S7). Furthermore, the module predominantly encompasses pathways that are broadly expressed over time, indicating that their presence is not exclusive to this module or its associated diatom assemblage.

Eigengene expression of modules M3, M8-M10, is correlated with the absolute abundance of a diverse range of genera, from copepods to diatoms. These modules encompass KOs from a diverse suite of pathways linked to fundamental aspects of (multicellular) cell growth, regulation, and maintenance such as ribosome synthesis and mRNA surveillance, signalling (based on transcripts homologous to genes involved in e.g. the ErbB and Hippo signalling cascades in mammals) and synthesis of glycolipids and glycosaminoglycans, vital for cell adhesion and communication and as structural components. Moreover, the presence of pathways associated with responses to bacterial and viral infections suggests an important role of host defence (Fig. S8, S9, S10, S11; Supplementary Dataset 2 & 3).

Module M6 eigengene expression, peaking in late spring and summer, correlates with the presence of dinoflagellate genera such as *Noctiluca*, *Tripos*, and *Effrenium*. Module M6 is linked to the synthesis of acarbose and validamycin, suggesting a capacity for antimicrobial compound production. Furthermore, module M6 is associated with the degradation of various aromatic compounds, emphasising the dinoflagellates’ role in organic matter breakdown and heterotrophic activities. Particularly noteworthy is the involvement of pathways related to the degradation of benzoate, flavonoids, and limonene, indicating the potential recycling of xenobiotic metabolites in sea surface waters by these assemblages. Additionally, module M6 is associated with metabolic functions encompassing vitamin B6 synthesis, polyketide sugar unit biosynthesis, biosynthesis of various plant secondary metabolites, and teichoic acid biosynthesis. Transcripts involved in these pathways were confirmed to be expressed by *Noctiluca* (Fig. S9).

Module M7 eigengene expression was strongly correlated with increased absolute abundances of the comb jelly genus *Mnemiopsis* and peaked in all stations (except 780) in August. This module is associated with homologs of mammalian genes involved in the ErbB signalling pathway, natural killer cell mediated cytotoxicity, and bacterial invasion of epithelial cells (Supplementary Dataset 2). While these pathways are detected across all months, a large part of the expression of these pathways in August could be attributed to *Mnemiopsis* (Fig. S8). Enzymes within these pathways are likely involved in either defence against microorganisms or the mechanisms of predation deployed by *Mnemiopsis*.

Lastly, module M4, with peak eigengene expression in summer, was associated with the degradation of limonene and flavonoids. Module M4’s eigengene expression showed a correlation with the abundance of the dinoflagellate genus *Tripos* and the cnidarian genus *Clytia*. Transcripts involved in these pathways were confirmed to be expressed by *Tripos* (Fig. S8) and *Noctiluca* (Fig. S9) with expression profiles peaking in summer, even though absolute abundances of *Noctiluca* did not significantly correlate with M4 eigengene expression (Fig. 4, Supplementary Dataset 3). Both genera are thus likely capable of catabolizing biotic carbon sources. Modules and their contents can be further explored in supplementary data (Supplementary Dataset 2).

### The case of dinoflagellates, diatoms, and *Phaeocystis*

We zoomed in on the most abundant taxonomic groups to gain a better insight into how functional richness links to the number of active species and how this relates to their ecology. The overall most abundant microeukaryote groups detected in the metatranscriptome were diatoms, dinoflagellates and *Phaeocystis* (Fig. 2).

From September to December, we observed higher abundances of diatom transcripts (Fig. S2 & S3) accompanied by a significant increase in the number of active diatom species (Fig. S12; Kruskal-Wallis, H(10,51) = 38.003, p = 3.791 *10^-5^). The fall assemblage was dominated by the genera *Trieres*, *Ditylum*, *Thalassiosira*, *Odontella*, and *Helicotheca*. A smaller peak in diatom abundance was observed at stations ZGO2 and 330 during June and July 2021, where the community was mainly comprised of *Guinardia* (based on EukProt). Stations closest to the Scheldt estuary, i.e. 700 and 780, exhibited the lowest diatom abundances. Diatom communities showed clear temporal structuring, aligning with temperature, and spatial gradients, including variations in nutrients, salinity, and SPM levels (Fig. S12d). For example, the eigengene expression of module M9, a set of KOs that correlates with the abundance of a *Thalassiosira* dominated diatom community, correlates significantly with elevated nutrient levels (Fig. 4 & Fig. S13; NO_3_^-^ r(10) = 0.41, p = 0.001; PO_4_^3-^ r(10) = 0.51, p = 3*10^-5^, Si r(10) 0.3, p = 0.02). The diatom genera that correlate most with module M9 eigengene expression are most abundant at station 130, 120, 330, and ZG02 (Fig. S2).

The diatom community’s functional richness appeared relatively consistent throughout the year and scales with the number of species present [Supplementary Fig. 19b, (linear model: MSE = 236,859.62 (SD = 103,071.89), R^2^ = 0.3044 (SD = 0.1924; polynomial model: MSE = 279,665.89 (SD = 130,398.46), R^2^ = 0.1720 (SD = 0.3299), a paired t-test revealed no statistically significant difference between the performance of the two models, p = 0.466]. In the fall, diatoms contributed most to the metabolic activity of the microeukaryotic plankton, exhibiting the highest expression of transcripts involved in primary production processes such as photosynthesis, carbon fixation, and fatty acid biosynthesis (Fig. 5 & Fig. S14). Predictions from the trophic mode classifier model by Lambert et al.^33^ consistently identified diatom species bins as autotrophic (Fig. S15 & S16, see Methods), although predictions were not feasible for smaller transcriptome bins.

**Figure 5.**
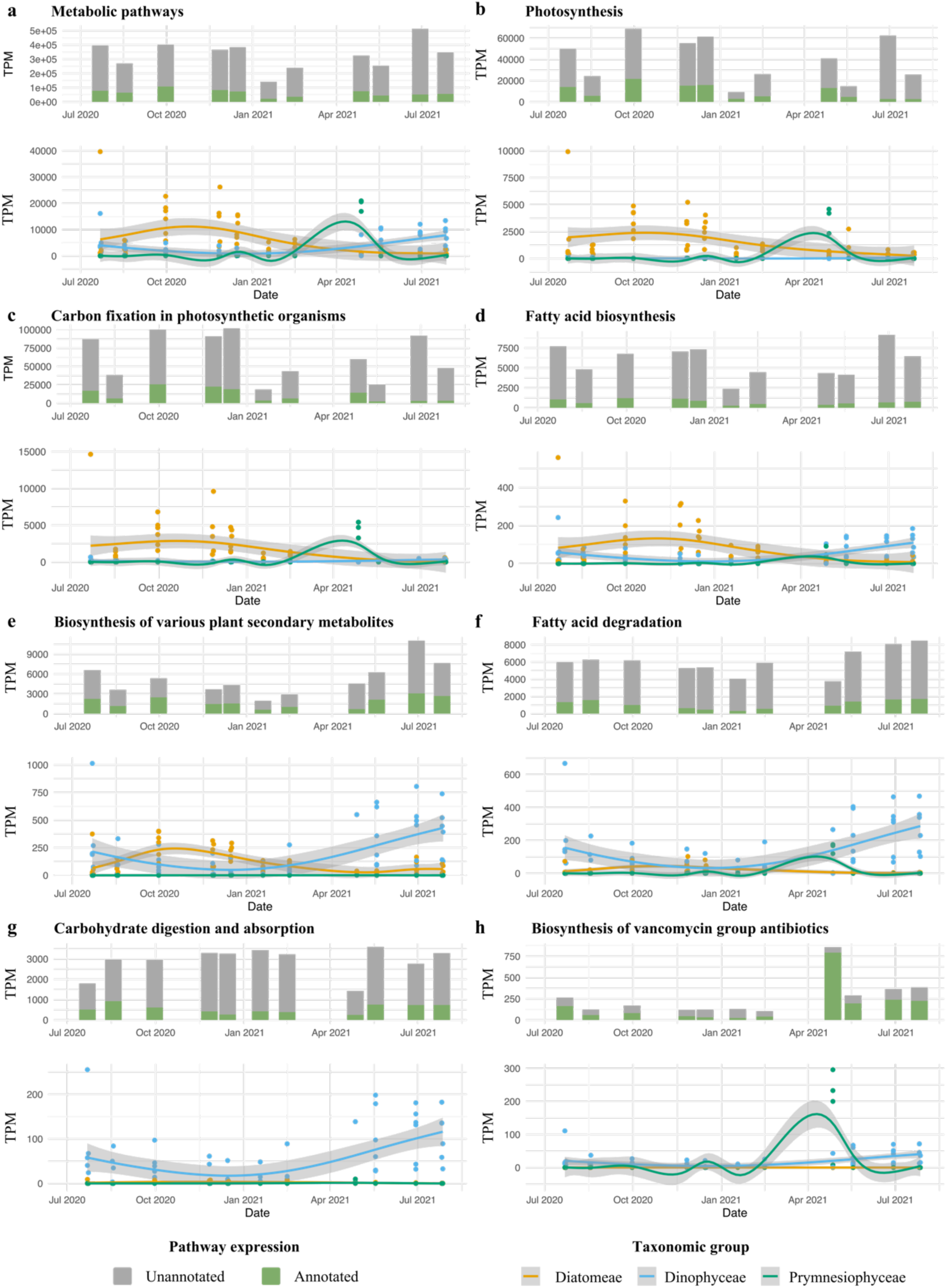
Expression of selected pathways and GAM regression fits for the expression of diatom, dinoflagellate, or *Phaeocystis* transcripts. Transcript per million (TPM) expression profiling of all transcripts associated with a given KEGG pathway of interest, alongside Generalised Additive Model (GAM) regression fits depicting the expression patterns of transcripts categorised within the pathways concerned that were annotated as diatoms, dinoflagellates, or *Phaeocystis* (>90% sequence identity) over time. The top figure in a panel contrasts the TPM expression levels of taxonomically annotated versus unannotated transcripts associated with the KEGG pathway. 8 pathways are displayed: **(a)** ‘metabolic pathways’, an umbrella term containing all metabolic activity, **(b)** photosynthesis, **(c)** carbon fixation in photosynthetic organisms, **(d)** fatty acid biosynthesis, **(e)** biosynthesis of various plant secondary metabolites, **(f)** fatty acid degradation, **(g)** carbohydrate digestion and absorption, and **(h)** biosynthesis of vancomycin antibiotics.

Dinoflagellates showed peak abundance from May to July, with *Noctiluca* being the most abundant genus (Fig. S17 & S18). The functional profiles of dinoflagellates varied predominantly along a temperature gradient (Fig. S12h). Module 6, associated with *Noctiluca*, also positively correlated with temperature (Fig. 4 & Fig. S13; r = 0.27, df = 10, p = 0.03), underscoring a seasonal trend in dinoflagellate abundance and activity. Notably, *Noctiluca* persisted in the colder, nutrient-rich, and less saline waters of stations 780 and 700 during fall and winter months (Fig. S17 & S18). Few active dinoflagellate species were detected across samples (Fig. S12). However, one to three active species of dinoflagellates can display a wide variety of functions, with maximum values that exceed double the number of diatom functional richness. Intriguingly, the Lambert et al. model^33^ predicted the EukProt-annotated *Noctiluca scintillans* taxonomic bin as consistently phototrophic (Fig. S15), while the PhyloDB annotated *Noctiluca scintillans* bin was predicted to be heterotrophic during the summer months, from May to September, and phototrophic in other months (Fig. S16). Similarly, based on PhyloDB annotations, *Tripos fusus* was predicted as heterotrophic in July 2021 and phototrophic during other months (Fig. S16), whereas the EukProt annotated *Tripos* bin was consistently predicted as phototrophic (Fig. S15). Very few transcripts involved in photosynthesis were linked to Noctiluca, while they do express transcripts involved in autophagy, fatty acid degradation, carbohydrate digestion and absorption, among others (Fig. 5, Fig. S14, Supplementary Dataset 3). The observed dinoflagellates’ ability to rely on both photo- and heterotrophy may contribute to the higher functional richness observed in dinoflagellate transcriptomes.

The prymnesiophyte *Phaeocystis globosa* bloomed in April, dominating the microplankton community in the Southern Bight in both abundance and metabolic activity (Fig. 2). The spring *Phaeocystis* bloom did not significantly correlate with distinct temperature, salinity, or nutrient values (Fig. S13). The Lambert et al.^33^ trophic model predictions based on both PhyloDB and EukProt annotation sources indicated that these *Phaeocystis* blooms were heterotrophic. They however also expressed transcripts involved in photosynthesis and carbon fixation (Fig. 5). Furthermore, we found that *Phaeocystis* was a major contributor to the expression of genes involved in specific pathways, such as the α-lipoic acid metabolism, the biosynthesis of polyketide sugar units, and the biosynthesis of antibiotics (Fig. 5 & Fig. S14). Surprisingly, for samples that contained *Phaeocystis* blooms we found very high expression of genes involved in certain pathways that were not annotated as *Phaeocystis*, for example zeatin biosynthesis or flagellar assembly. These might constitute vital processes in the *Phaeocystis* life cycle that are currently not represented in the reference transcriptome, for example an investment in motile microflagellate cells after the blooming phase^34^ (Fig. S14).

At the time of year when diatoms, dinoflagellates or *Phaeocystis* are most abundant, this is often accompanied by a high expression of genes related to viral infection (Fig. S14). This likely suggests a heightened viral activity or prevalence, as indicated by the elevated expression of homologs of genes involved in pathways related to infections of coronaviruses and papillomaviruses in humans. This suggest viral infections may be a significant factor in microeukaryotic population dynamics^35^.

## Discussion

The observed seasonal shifts in taxonomic assignments of gene transcripts echo well-documented phenological patterns in the Southern Bight. These include the early spring bloom of *Phaeocystis globosa*^36^, the prevalence of *Noctiluca scintillans* between May and July^37^, and the late summer bloom of the invasive ctenophore *Mnemiopsis leidyi*^38^. These phenological patterns are intricately linked to the seasonal oscillations of abiotic factors such as temperature, nutrient availability, or solar irradiance. Our study further reveals seasonal partitioning of key metabolic pathways among the predominant groups. For example, we observed a shift between diatoms and dinoflagellates alternating in their contribution to the production of fatty acids and other secondary metabolites.

The first plankton bloom of the year in the Southern Bight occurs in diatoms in spring, leveraging the elevated temperatures and sunlight to utilise the accumulated winter nutrients derived from river discharge, atmospheric deposition, and sediment resuspension^28,39^. We observed a rise in metabolic activity of diatoms already in February, preceding the actual increase in cell densities detected by imaging in spring. After the first diatom bloom, *Phaeocystis globosa* capitalises on favourable light and nutrient concentrations and forms colonies that heavily invest in primary production^40,41^. Recent laboratory studies have shown that these *Phaeocystis* blooms attract a specialised bacterial community^54^. Our *in-situ* findings suggest that *Phaeocystis* implements a heterotrophic strategy at later stages of the bloom. This might represent an adaptive strategy, where *Phaeocystis* cultivates and attracts bacteria during favourable periods and subsequently feeds on them following bloom senescence and nutrient depletion. While we did not find clear evidence of mixotrophy for *Phaeocystis* in trophic mode predictions, we observed the expression of transcripts involved in both photosynthesis and heterotrophy^41^.

*Noctiluca scintillans* likely deploys different feeding strategies across seasons, potentially underpinning their ecological success. The pronounced functional richness of dinoflagellates confirms an elevated level of metabolic flexibility, a hypothesis supported by the ambiguous trophic mode predictions and the occurrence of degradation, digestion, and absorption pathways. This versatility potentially confers a competitive edge in fluctuating environments^42^.

In fall and early winter, diatoms exhibited higher expression of transcripts involved in primary production processes such as photosynthesis, carbon fixation, and fatty acid biosynthesis. This diatom proliferation in the fall deviates from historical records, suggesting possible shifts in ecological processes or broader environmental changes^5,43,44^.

The increased expression of antimicrobial or immunity-related pathways during blooms of *Phaeocystis globosa* and *Mnemiopsis leidyi* points towards the critical role of biotic interactions in shaping community dynamics^45^. The occurrence of transcripts associated with the production of toxins in *Phaeocystis* suggests a mechanism by which the *Phaeocystis* colonies could defend against a *Hemistasia* infection^29^. These observations align with the growing consensus on the importance of parasites and pathogens in regulating bloom events^46^.

### Conclusion and perspective

This study utilised metatranscriptomics to reveal phenological patterns and functional properties of microeukaryotes in the Southern Bight. Our findings demonstrate the potential of longitudinal metatranscriptomic time series in enhancing our understanding of microbial dynamics. The dataset presented supports future research aimed at developing predictive models for ecosystem behaviour^47,48^. Continued exploration of these crucial microbial communities is vital, especially given the accelerating pace of climatic and environmental shifts.

## Methods

### Sampling

The data presented in this study was obtained from samples collected on campaigns from July 2020 to July 2021. Several stations in the Belgian North Sea were sampled every month with the R/V Simon Stevin^49,50^ (offshore stations: ZG02, 330, 700; nearshore stations: 120, 130, 700, see Fig. 1 and Supplementary Dataset 1). When at sea, navigation data, meteorological and oceanographic data were measured using the ship’s underway system. At each sampling station, temperature (°C) and depth (m) profile (CTD) data were collected with a Seabird SBE25plus CTD (Sea-Bird Scientific, Bellevue, WA USA). Additionally, the following parameters were quantified: salinity of the water body, expressed in practical salinity units (PSU) and the concentration of suspended particulate matter (SPM) in the water body (expressed in mg/L). Water samples were taken with Niskin bottles at a depth of 3 m and were analysed for nutrient and pigment concentrations. For more detail on sampling procedures, see Mortelmans et al. (2019).

Microphytoplankton biomass was estimated using FlowCam analysis, which involved collecting 50 L of sea surface water, filtering, and processing with a FlowCam VS-4 benchtop model (Fluid Imaging Technologies, Yarmouth, Maine, USA.) to classify particles into distinct taxonomic groups; further details are provided in the supplementary methods and Martínez et al. (2020)^11^.

Samples for metatranscriptomic analysis were collected as follows: at each sampling location, 50 L of sea surface water was manually collected with stainless-steel buckets and filtered through two stacked stainless-steel sieves with mesh sizes of 250 and 50 µm. In case of extreme bloom events, smaller volumes were filtered due to the high biomass clogging the filters. From the 50 µm sieve the collected residue was resuspended in 9 - 45 mL of seawater, with the eluate volume depending on the residue biomass (Supplementary Dataset 1). The samples were then stored in cryovials and flash-frozen in liquid nitrogen within five minutes of collection. Back in port, samples were transferred to a -80 °C ultra-freezer until RNA extraction.

### Extraction & quality control of RNA, library preparation & sequencing of cDNA

RNA was extracted with the RNeasy Mini Kit (Qiagen), according to a modified version of the manufacturer’s protocol. Briefly, samples were thawed, centrifuged, and the supernatant seawater was removed. Lysis buffer with 700 mg of RNA-free silica beads (350 mg each of 0.1 and 0.5mm) was added to the sample after which samples underwent two cycles of homogenization and cooling on a metal cooling block (−20 °C) for one minute each. The RNA was then extracted using the kit’s spin columns. Depending on the season and biomass of microplankton, different starting volumes were used for RNA extraction (Supplementary Dataset 1). The extracted RNA was eluted in 52 µL of RNAse-free water and stored in the -80 °C freezer. 2 µL of the total RNA eluate was used for quality and yield analysis. Total RNA yield was measured using a Qubit fluorometer (Invitrogen), RNA quality was assessed using a Bioanalyzer (Agilent). After extraction, samples were shipped to the VIB Nucleomics core facility (https://nucleomicscore.sites.vib.be/) on dry ice. There, RNA concentration and purity were determined spectrophotometrically using the Nanodrop ND-8000 (Nanodrop Technologies) and RNA integrity was assessed using a Bioanalyzer 2100 (Agilent). A standard volume of ERCC spike-in Mix was added before starting the library preparation. Per sample, an amount of 200 ng of total RNA was used as input. Using the Illumina TruSeq® Stranded mRNA Sample Prep Kit (protocol version: # 1000000040498 v00 October 2017), poly-A containing mRNA molecules were purified from the total RNA input using poly-T oligo-attached magnetic beads. In a reverse transcription reaction using random primers, RNA was converted into first strand cDNA and subsequently converted into double-stranded cDNA in a second strand cDNA synthesis reaction using DNA Polymerase I and RNAse H. The cDNA fragments were extended with a single ‘A’ base to the 3’ ends of the blunt-ended cDNA fragments after which multiple indexing adapters were ligated introducing different barcodes for each sample. Finally, enrichment PCR was carried out to enrich those DNA fragments that have adapter molecules on both ends and to amplify the amount of DNA in the library. Sequence libraries of each sample were equimolarly pooled and sequenced on Illumina NovaSeq 6000 (SP300 flow cell, PE150 (151-8-8-151)) at the VIB Nucleomics Core.

### Preprocessing, assembly, and translation

Raw sequences were inspected using FastQC and MultiQC^51^. Sequencing adapters were trimmed from the reads using Trimmomatic^52^ (version 0.39, parameters: ILLUMINACLIP:ADAPTERS:2:30:7,LEADING:2,TRAILING:2,SLIDINGWINDOW:4:2, MINLEN:50). Ribosomal RNA sequences were removed using RiboDetector^53^. Quality-controlled, trimmed non-rRNA PE reads were assembled in nucleotide space using rnaSPAdes (version 3.15.3, parameters: --rna -k 75,99,127)^54^. All assemblies were then combined, and clustered at 95% identity using MMseqs2 easy-linclust algorithm (version 13-45111)^55,56^. The contig names of the resulting metatranscriptome were standardised with anvi-script-reformat-fasta from Anvio (version 7.1) to reduce their size and facilitate subsequent analysis^57^. Assembled transcripts that matched the ERCC92 spike-in RNA sequences were identified using MMseqs2 easy-search and removed, resulting in the final metatranscriptome^56^. Protein coding regions in the *de novo* assembled transcripts were identified by Transdecoder (version 5.5.0, parameters: --single-best-only)^58^.

### Transcript annotation and quantification

To assign taxonomic information to the assembled and translated sequences, they were searched against EukProt^20^ and an extended version of PhyloDB (version 1.075; https://allenlab.ucsd.edu/data/), using MMseqs2^56^. The PhyloDB database was extended with sequence data from ENA for species that were identified in FlowCam data but were not included in the database^59^. If a published shotgun assembly transcriptome existed for that species, it was added to the reference database. For both databases, best alignment hits with a sequence identity > 90% or > 60% (>60% sequence identity when considering taxonomic classes) were retained for subsequent analysis. Peptide translations of the *de novo* assembly were functionally annotated using the eggNOG reference database^60^ and the eggNOG-mapper tool (version 2.1.7)^61^. Transcript and ERCC spike-in standards were quantified using Kallisto, yielding transcript per million (TPM) counts for every transcript^62^. Relative abundance of a given taxonomic group was calculated as the sum of all TPM values belonging to that level divided by the total TPM sum of all taxonomically annotated transcripts for that sample or month, ignoring unannotated transcripts. Transcripts per L were calculated by relating the TPM counts to the spike-in recovery values while accounting for sample processing volumes^26^ (see Supplementary Methods).

### Statistical analysis

Principal component analyses were performed on pre-processed log-transformed transcripts using sci-kit learn in python (version 1.1.3)^64,65^. Arrows representing the correlation between the first two principal components and environmental parameters were added to the plots.

A weighted gene co-expression network analysis was performed using the WGCNA package^31^ (version 1.72-1) in R^66^. The approach was modified from the analysis pipeline described in Cohen et al, 2021^42^. Briefly, we aimed to identify co-expression modules of Kyoto Encyclopedia of Genes and Genomes (KEGG) KO identifiers from log-normalised count sums of KEGG KOs over annotated transcripts (see Supplementary Methods). We calculated module eigengenes and assessed their correlations with environmental parameters and taxonomic groups’ absolute TPM abundances. To identify metabolic pathways associated with a given module eigengene, we performed Mann-Whitney U tests to assess which pathways were more represented at the top of a ranked list of KEGG KO identifiers, ordered according to decreasing module membership, than expected by chance. Module membership of a pathway was defined as the Pearson correlation between module eigengene expression and KEGG KO identifier. Resulting p-values were adjusted for multiple testing with the Benjamini-Hochberg procedure, which controls the false discovery rate^67^.

KEGG pathway expression was analysed by collecting the total expression of assembled transcripts that were annotated with a KO involved in that pathway. For these, we retrieved EukProt annotations with >90 % sequence identity to be able to calculate the annotated and unannotated fraction, or to assess which taxonomic group contributed most to the given pathway. For pathways identified as characteristic of modules identified through WGCNA, we visualised the total TPM expression of transcripts involved in each pathway and the fraction that could be assigned to the genus whose absolute abundances correlated most with expression of the module eigengene. Generalised additive models (GAMs) were constructed with the mgcv package in R (version 1.8-40) to assess the temporal dynamics of gene expression for KEGG pathways^66,68^. To complement the GAM analysis, we aggregated the data by month and calculated the total TPM for each pathway, both for all transcripts and transcripts that had a EukProt taxonomic annotation (>90% sequence identity). This was visualised alongside the GAM fits to provide a more comprehensive view of the pathway activity over time^69^.

To obtain active species richness estimates of samples, taxonomic bins of transcripts with > 90% sequence identity with sequences in reference databases were considered. Only taxonomic bins that had more than 100 non-zero expressed transcripts in at least one sample were retained, and their occurrence per sample was quantified. Functional richness was defined as the amount of unique KEGG KO identifiers found in a sample.

We used the classification machine learning model by Lambert et al. (2022) to predict the trophic mode of protist taxonomic bins^33^. The training data, parameter settings and selected features were used as in the original study (we implemented the XGBoost classifier with reported precision of 88 ± 10% in the original publication). Taxonomically annotated transcripts were binned to species level (>90% seq. id), and the model was trained with the MinMax-scaled MMETSP training dataset from the original publication containing the union of selected features^33^. Predictions were then generated for each sample by subsetting each profile to contain the union of selected features, imputing missing values with zeros, and scaling profiles with the pre-fitted MinMax scaler. Predictions could only be made for subsetted transcriptional profiles that contained > 800 non-zero expressed PFAMs, as required by the Lambert et al. model^33^. Monthly consensus predictions were determined by majority vote of the month’s individual samples’ predictions.

## Acknowledgements

The authors are grateful to the Flanders Marine Institute (VLIZ) and the LifeWatch project, a Flemish contribution to the LifeWatch ESFRI by VLIZ, for their financial and technical support. A special thanks to the crew of the research vessel Simon Stevin. The computing resources and services used in this work were provided by the VSC (Flemish Supercomputer Center), funded by the Research Foundation - Flanders (FWO) and the Flemish Government.

## Author contributions

M.P. collected the samples alongside the LifeWatch campaigns organized by J.M. J.M. provided the environmental data, pigment data, and nutrient data. M.P. and P.H. developed the extraction protocol, M.P. performed the lab work supervised by P.H. M.P. performed bioinformatics and statistical analysis, guided by P.H. and S.M. R.L. did the FlowCam image processing, model training and evaluation, R.L. and M.P. jointly analysed the resulting FlowCam data. P.H. conceived the project, P.H. and S.M. supervised the project. M.P., S.M. and P.H. wrote the manuscript which was revised by all authors.

## Data availability

The metatranscriptomic data generated in this study is available on SRA under the BioProject PRJNA1021244. Analysis scripts are available on GitHub (https://github.com/MichielPerneel/BPNS_seasonal_MTX). All other data supporting the findings of this study are provided in Supplementary Data.

## Competing interests

The authors declare no competing interests.

## Supplementary Information

### Supplementary Figures

**Supplementary Figure 1.**
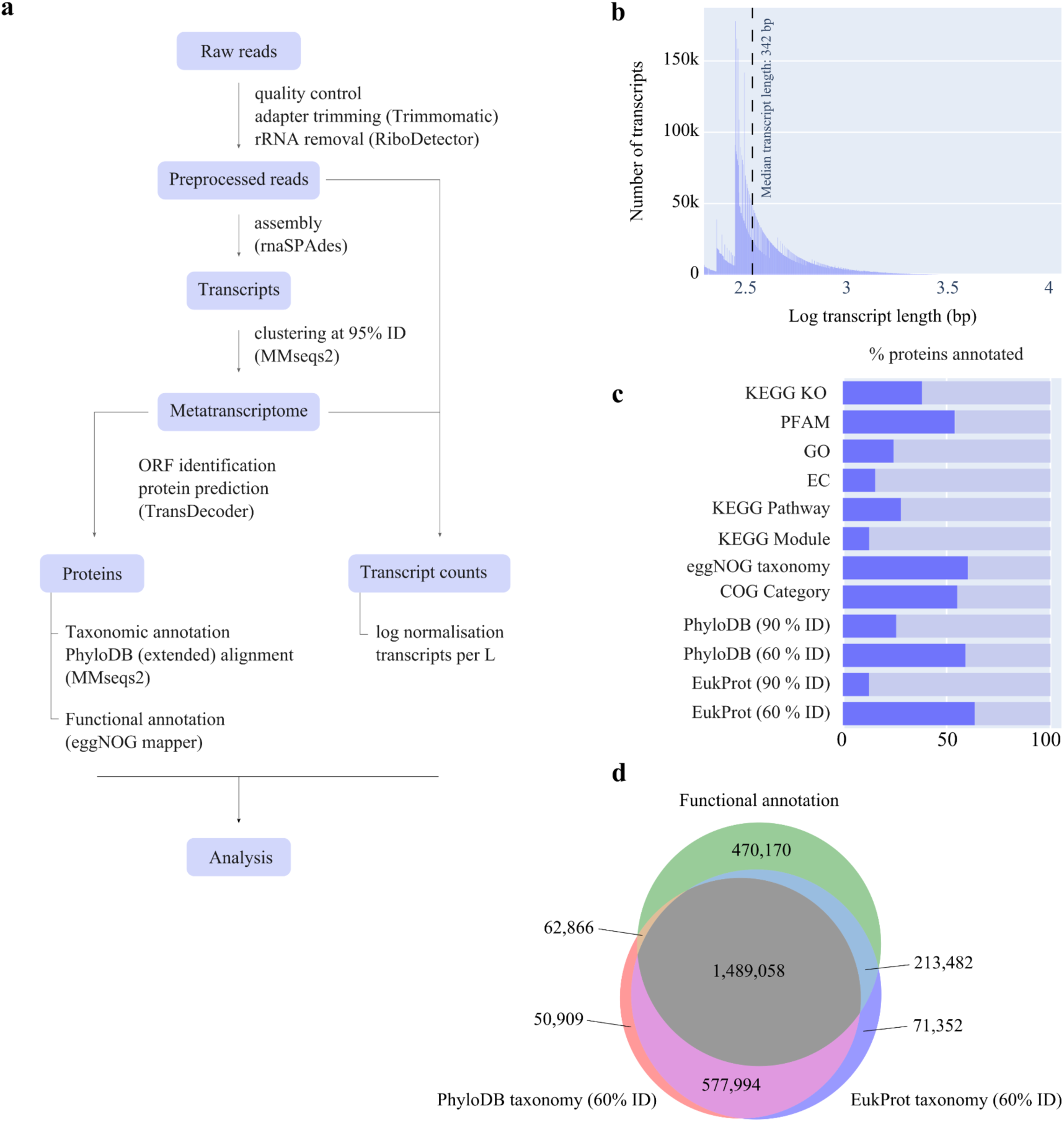
Bioinformatic workflow and metatranscriptome metrics. **a)** Bioinformatics workflow. **b)** Log transcript length distribution in the de novo assembled metatranscriptome. The dotted line indicates the median transcript length (342 bp). **c)** The percent of predicted proteins for which annotation information could be found using eggNOG mapper (resulting annotations consist of KEGG KOs (KEGG orthology identifiers), PFAM (protein families), GO (gene ontology categories), EC (Enzyme Commission numbers), KEGG Pathway identifiers, KEGG Module descriptions, eggNOG taxonomic classification, COG (clusters of orthologous groups)) and MMseqs2 alignment to both a custom extended version of PhyloDB and EukProt, both with a 60% and 90% sequence identity cut-off threshold value. **d)** Venn diagram showing the number of predicted proteins with a PhyloDB taxonomic annotation (>60% sequence identity with reference), and/or a EukProt taxonomic annotation (>60% sequence identity with reference), and/or functionally annotated predicted proteins with eggNOG functional information.

**Supplementary Figure 2.**
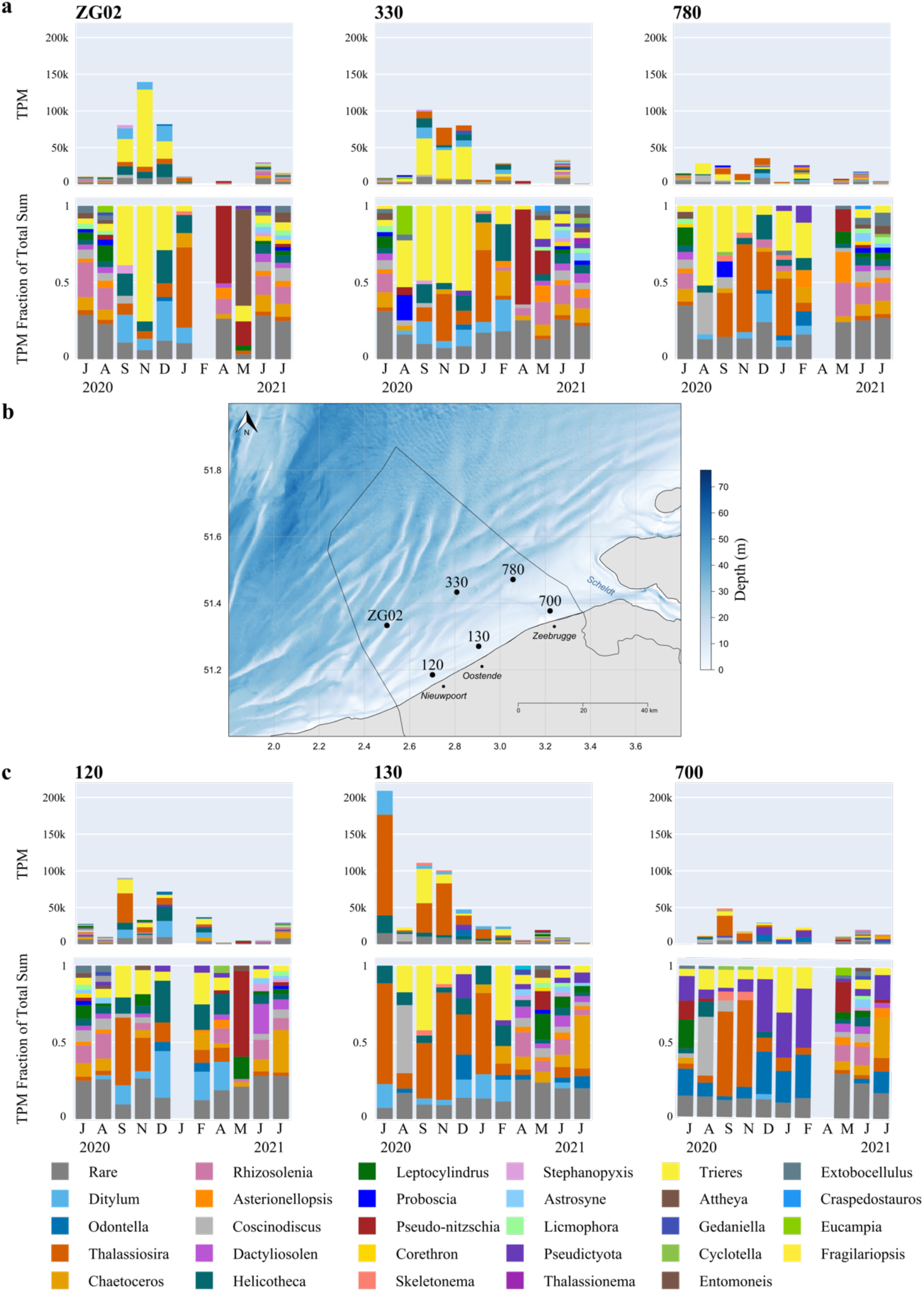
Monthly absolute and relative abundances of EukProt diatom genera at each sampling station. **a)** Monthly transcript abundance and relative distribution of diatom genera annotated using EukProt, per offshore sampling station. Transcript abundance represents the sum of transcripts per million (TPM) belonging to a taxonomic group. Relative abundance was calculated as the sum of TPM for that group for a given sample, divided by the total TPM of all genera found in that sample. When relative abundance of a genus was <2 %, they were labelled as ‘rare’. **b)** Spatial location of the 6 sampling stations in the Belgian North Sea. **c)** Monthly transcript abundance and relative distribution of diatom genera annotated using EukProt, per nearshore sampling station. Transcript abundance and relative abundance were calculated the same as for the offshore stations.

**Supplementary Figure 3.**
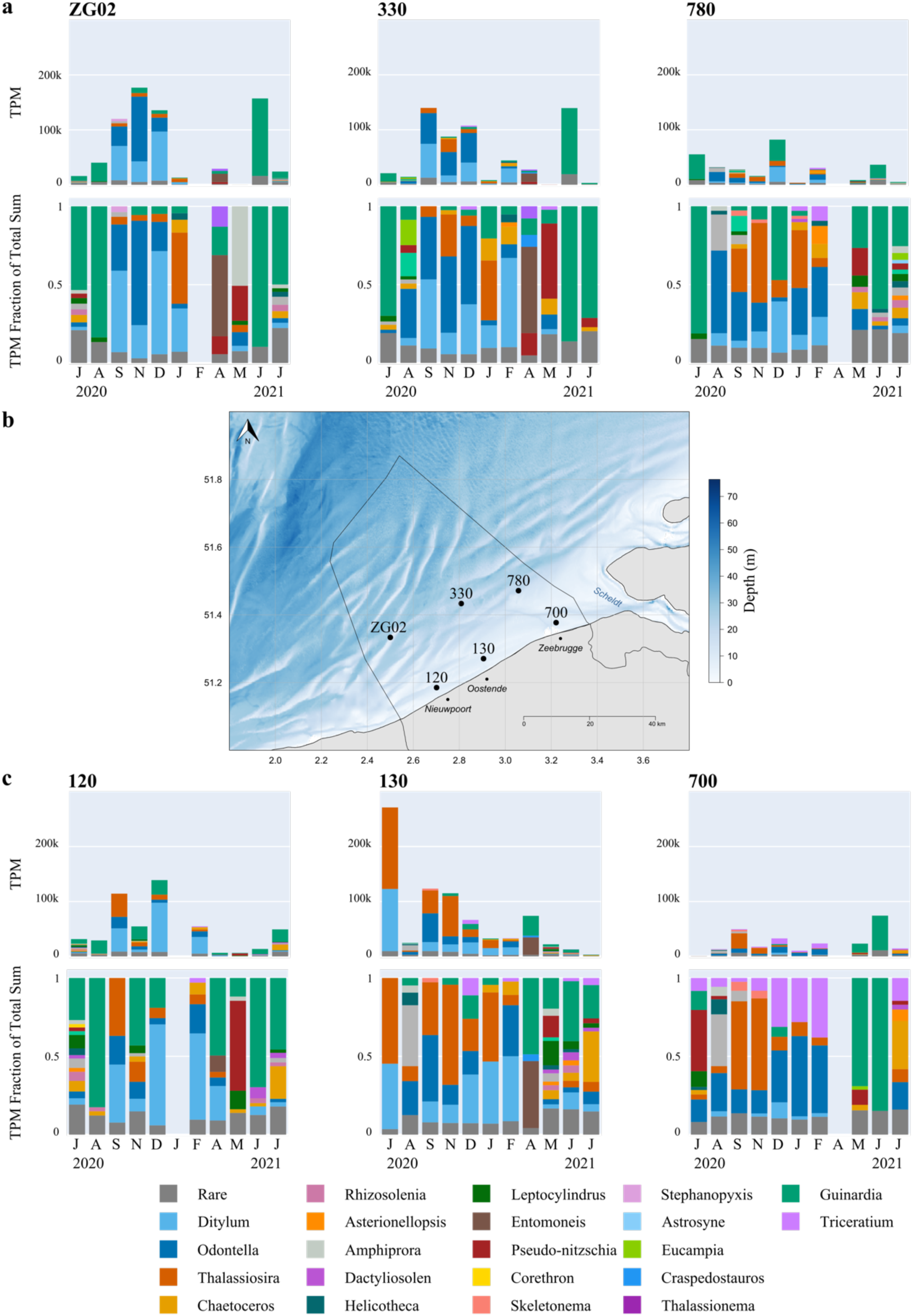
Monthly absolute and relative abundances of PhyloDB diatom genera at each sampling station. **a)** Monthly transcript abundance and relative distribution of diatom genera annotated using PhyloDB, per offshore sampling station. Transcript abundance represents the sum of transcripts per million (TPM) belonging to a taxonomic group. Relative abundance was calculated as the sum of TPM for that group for a given sample, divided by the total TPM of all genera found in that sample. When relative abundance of a genus was <2 %, they were labelled as ‘rare’. **b)** Spatial location of the 6 sampling stations in the Belgian North Sea. **c)** Monthly transcript abundance and relative distribution of diatom genera annotated using PhyloDB, per nearshore sampling station. Transcript abundance and relative abundance were calculated the same as for the offshore stations.

**Supplementary Figure 4.**
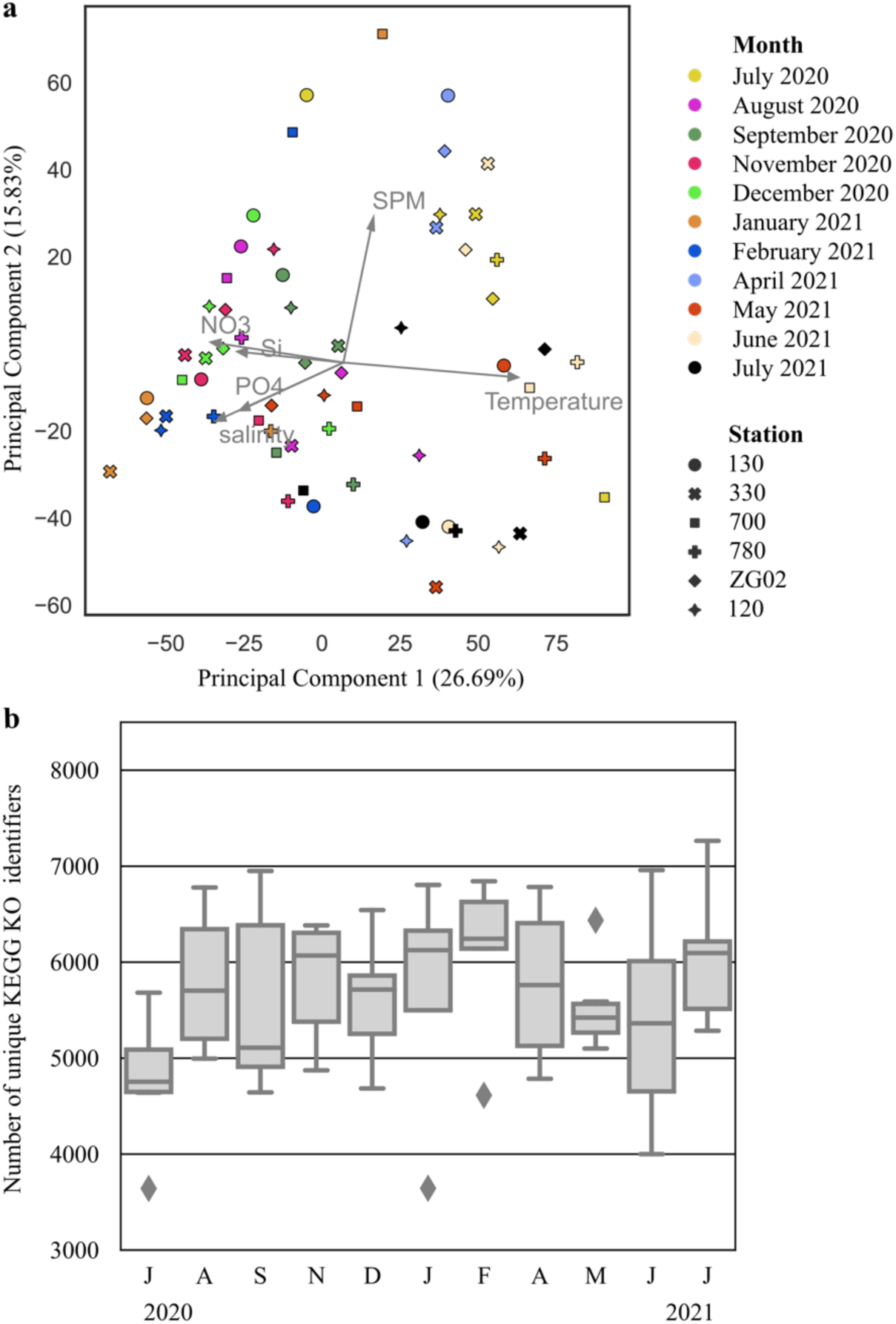
Seasonal changes in ecosystem functional richness and composition. **a)** Principal Component Analysis of log transformed transcript per million (TPM) expression data (summed per KO identifier per sample). Colours represent sampling months; shapes reflect sampling stations. Arrows indicate correlation of principal components with environmental parameters. **b)** Boxplot showing the number of unique KO identifiers detected per month per sample from July 2020 to July 2021. Boxes encompass the interquartile range, with the central line showing the median. Whiskers extend to the furthest data points within 1.5 times the interquartile range below the first or above the third quartile.

**Supplementary Figure 5.**
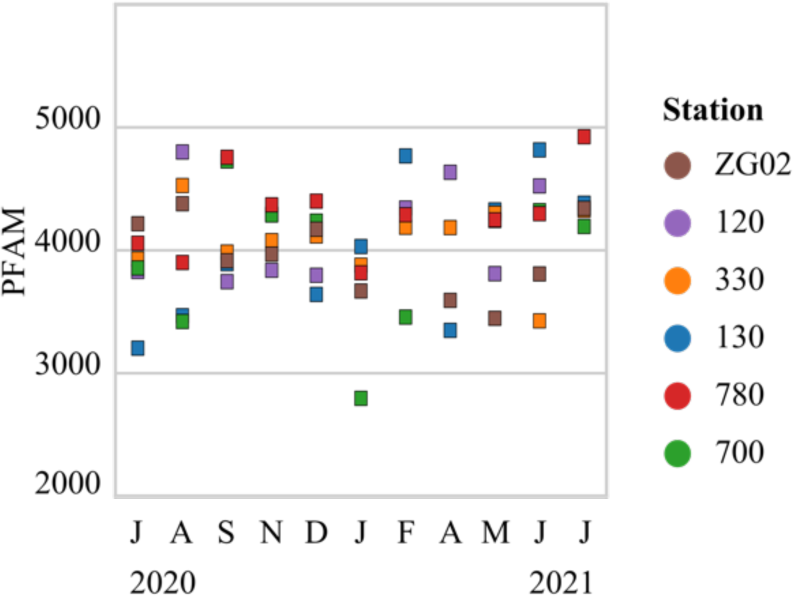
The number of unique PFAM families observed per sample per month, coloured according to sampling station.

**Supplementary Figure 6.**
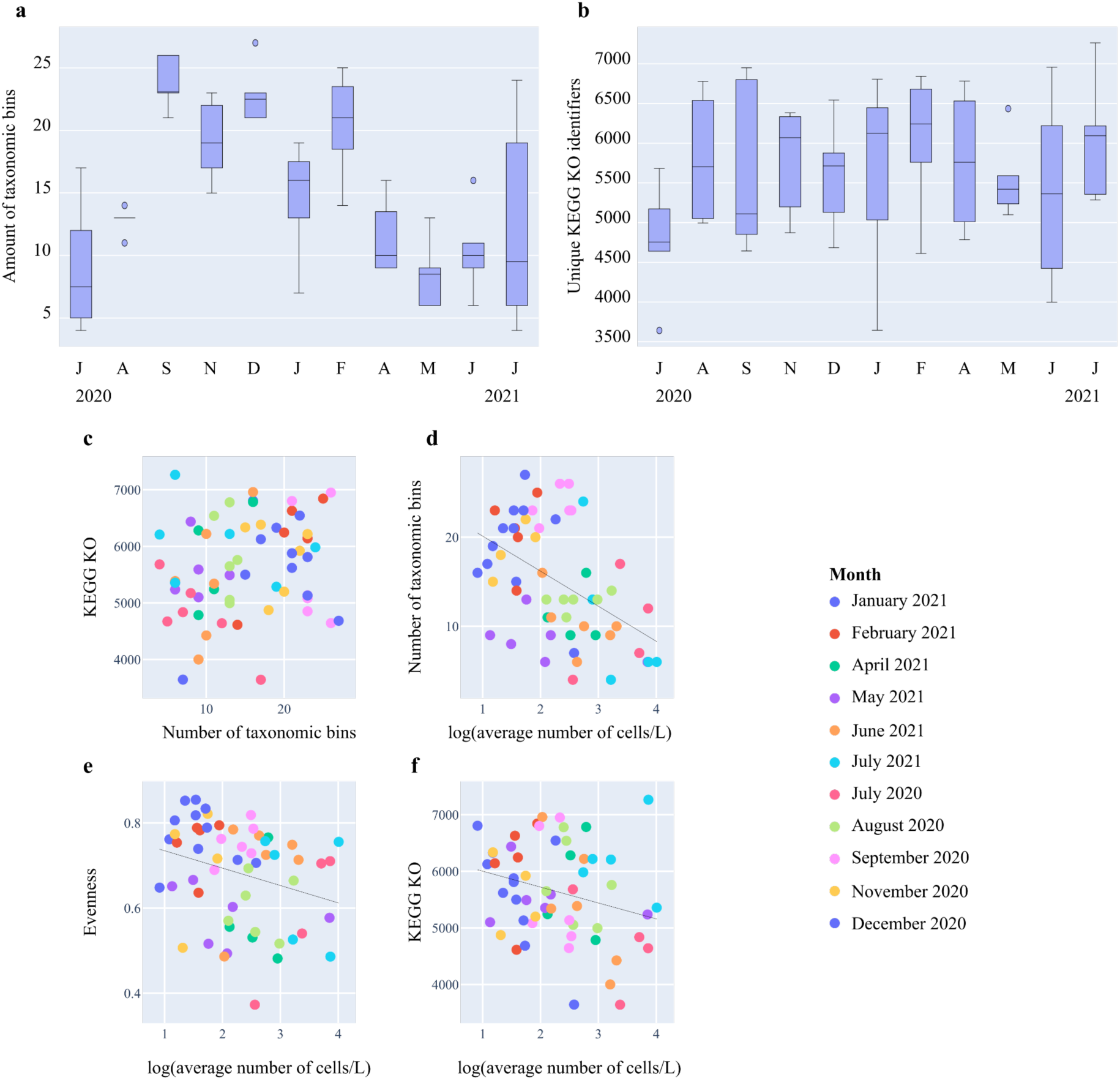
The relation between functional richness, species richness, and biomass. **a-b)** Boxplots of the number of active species (taxonomic species bins with a minimum of 100 transcripts with non-zero expression in at least one sample), and the number of unique KEGG KO identifiers in a sample over time. Boxes encompass interquartile range, with the central line showing median. Whiskers extend to the furthest data point within 1.5 times the interquartile range. **c)** The number of active species in relation to the number of unique KEGG identifiers across samples (r(60) = 0.18, p = 0.151). **d)** The log-transformed average estimate of cells per L of sea water in relation to the number of active species (r(55) = -0.48, p = 0.0002). **e)** The log-transformed average number of cells per L in relation to the evenness (Shannon diversity divided by the number of species) (r(55) = -0.28, p = 0.0364). **f)** The log-transformed average number of cells per L in relation to the number of unique KEGG identifiers across samples (r(55) = 0.26, p = 0.0492).

**Supplementary Figure 7.**
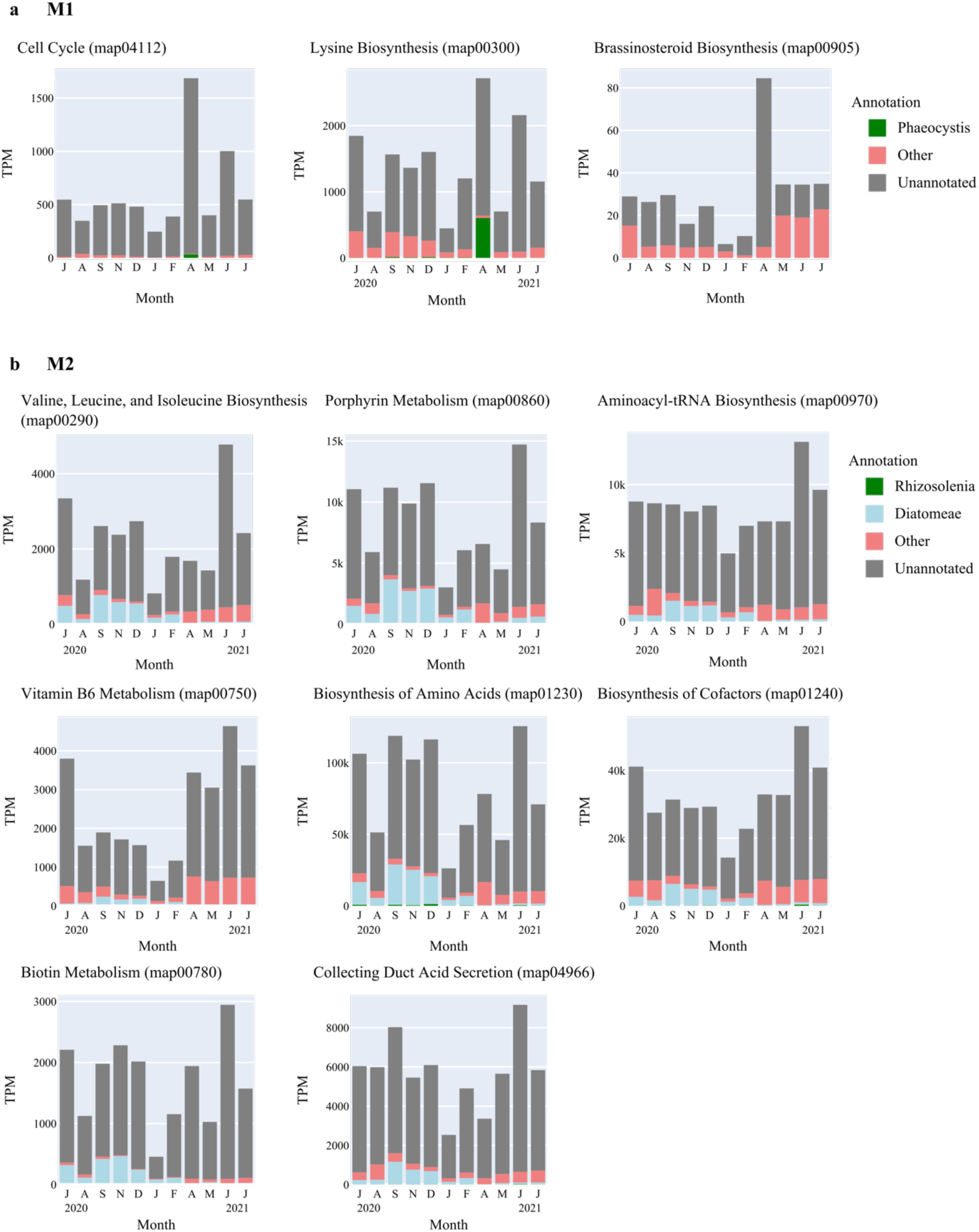
Expression of pathways characteristic of module 1 and 2. Each plot represents a characteristic pathway for module 1 (panel a) or module 2 (panel b), as determined by Mann-Whitney U tests assessing the correlation between the TPM expression levels of a pathways’ KO identifiers and the eigengenes of the respective module. Each plot shows the monthly TPM values for a characteristic pathway, incorporating all transcripts annotated to KOs involved in this pathway and summed across stations. Green bars indicate the fraction of transcript expression annotated to the genus whose absolute abundance profile is most highly correlated with the eigengene expression of the module concerned. Blue bars indicate the fraction of transcript expression annotated to the same taxonomic class as the genus represented by the green bars. Red bars indicate fraction of transcript expression annotated to other groups. Grey bars denote taxonomically unannotated transcripts.

**Supplementary Figure 8.**
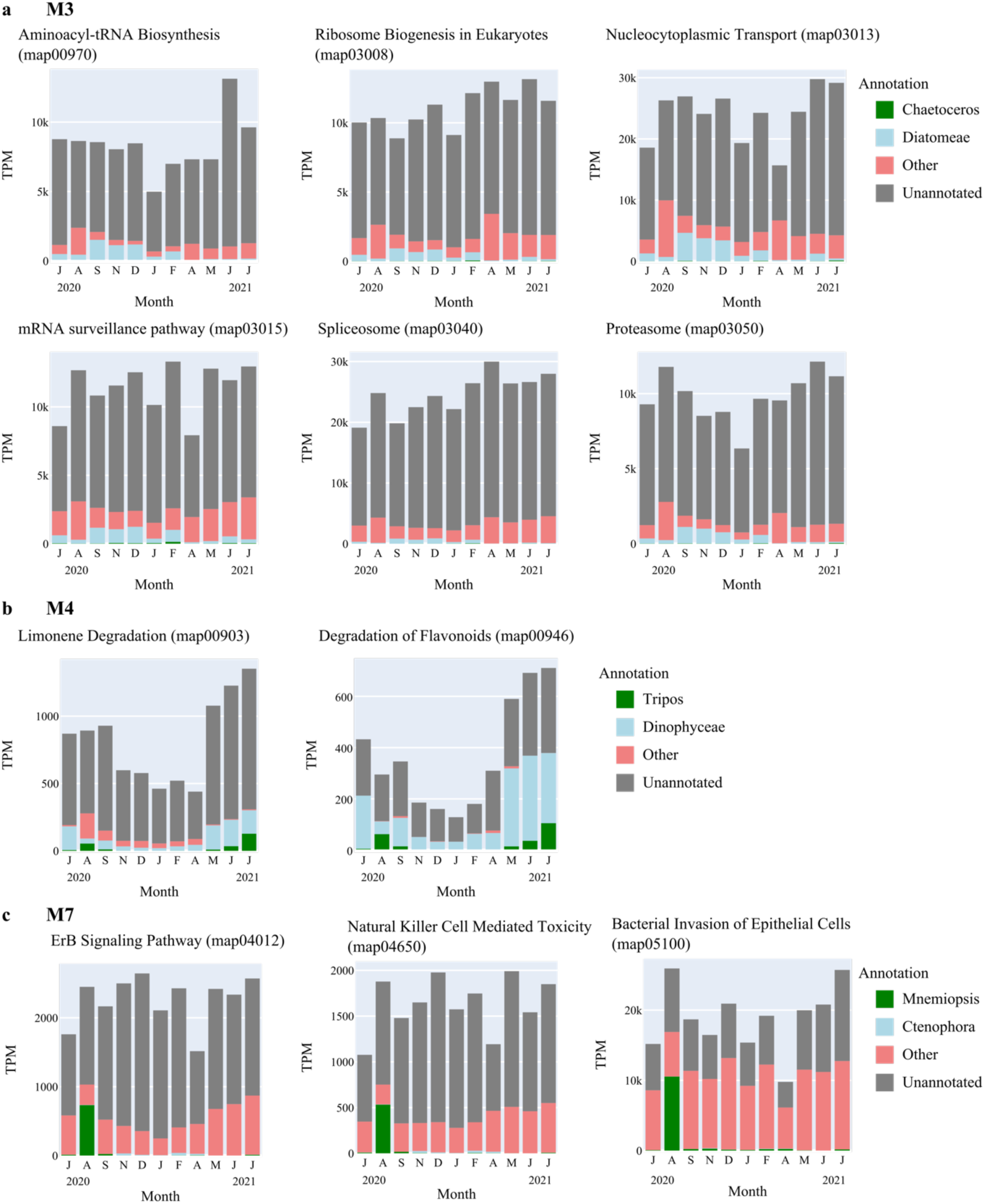
Expression of pathways characteristic of module 3, 4, and 7. Each plot represents a characteristic pathway for module 3 (panel a), module 4 (panel b) or module 7 (panel c) as determined by Mann-Whitney U tests assessing the correlation between the TPM expression levels of a pathways’ KO identifiers and the eigengenes of the respective module. Each plot shows the monthly TPM values for a characteristic pathway, incorporating all transcripts annotated to KOs involved in this pathway and summed across stations. Green bars indicate the fraction of transcript expression annotated to the genus whose absolute abundance profile is most highly correlated with the eigengene expression of the module concerned. Blue bars indicate the fraction of transcript expression annotated to the same taxonomic class as the genus represented by the green bars. Red bars indicate fraction of transcript expression annotated to other groups. Grey bars denote taxonomically unannotated transcripts.

**Supplementary Figure 9.**
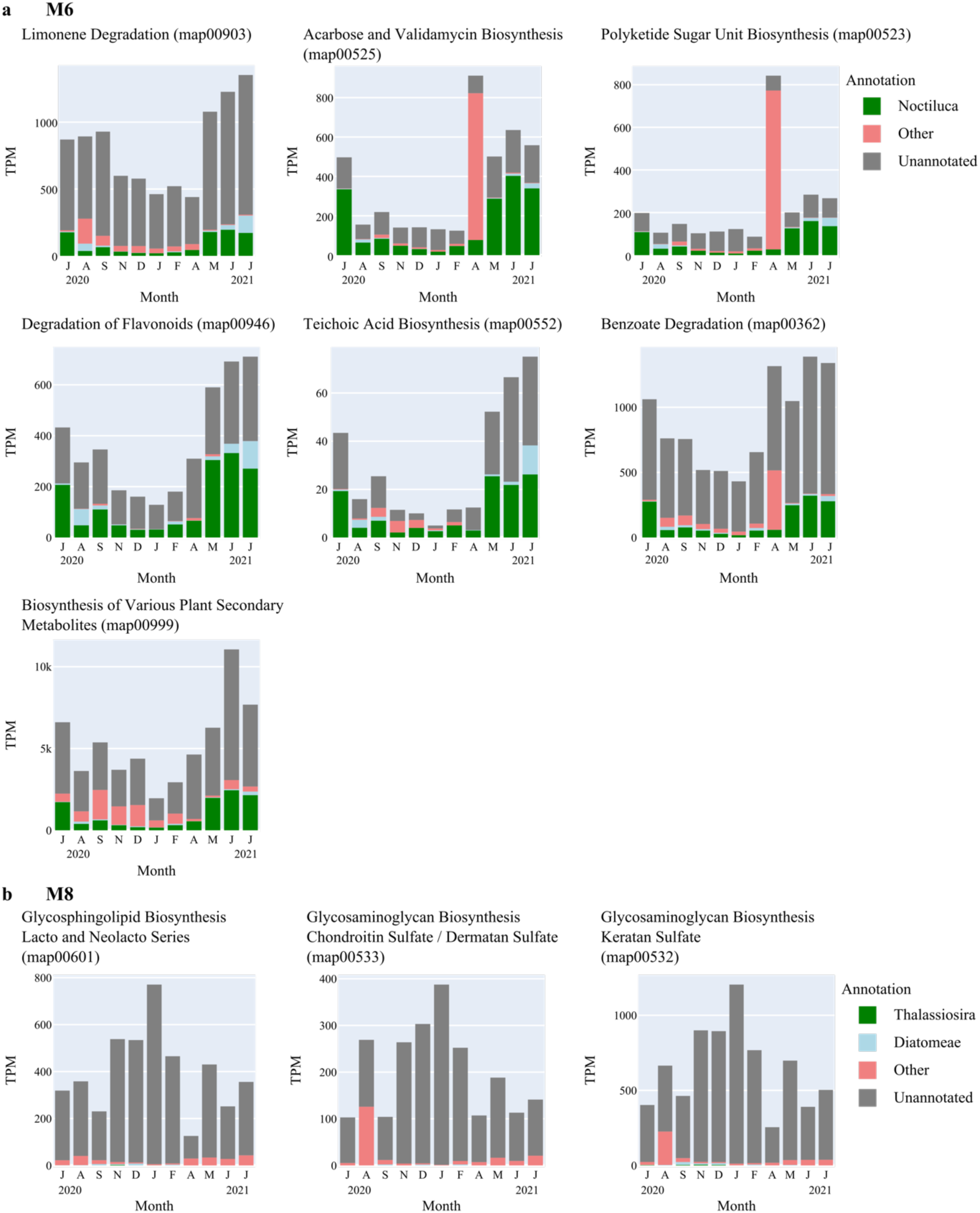
Expression of pathways characteristic of module 6 and 8. Each plot represents a characteristic pathway for module 6 (panel a) or module 8 (panel b), as determined by Mann-Whitney U tests assessing the correlation between the TPM expression levels of a pathways’ KO identifiers and the eigengenes of the respective module. Each plot shows the monthly TPM values for a characteristic pathway, incorporating all transcripts annotated to KOs involved in this pathway and summed across stations. Green bars indicate the fraction of transcript expression annotated to the genus whose absolute abundance profile is most highly correlated with the eigengene expression of the module concerned. Blue bars indicate the fraction of transcript expression annotated to the same taxonomic class as the genus represented by the green bars. Red bars indicate fraction of transcript expression annotated to other groups. Grey bars denote taxonomically unannotated transcripts.

**Supplementary Figure 10.**
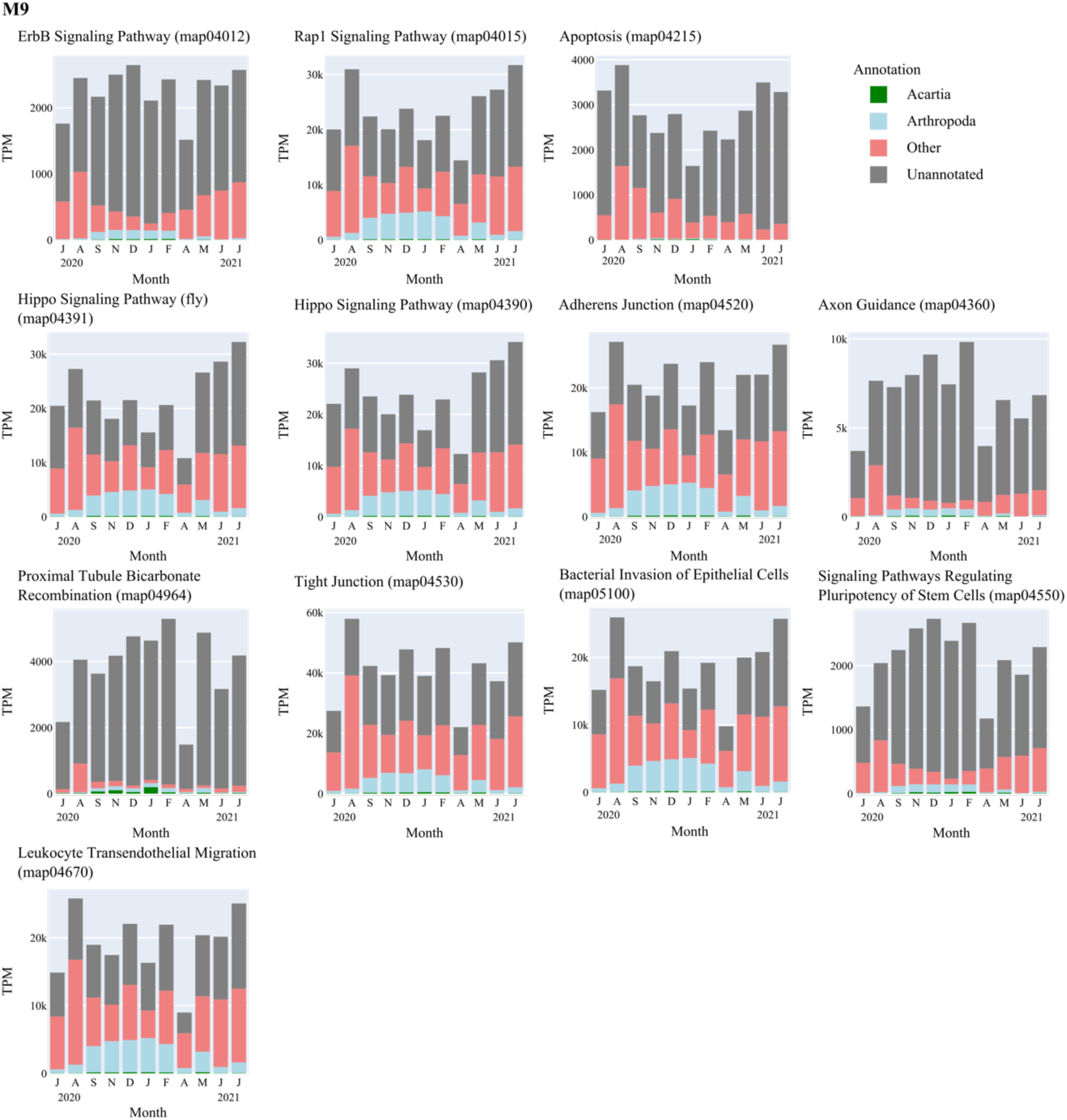
Expression of pathways characteristic of module 9. Characteristic pathways for module 9, as determined by Mann-Whitney U tests assessing the correlation between the TPM expression levels of a pathways’ KO identifiers and the eigengenes of the respective module. Each plot shows the monthly TPM values for a characteristic pathway, incorporating all transcripts annotated to KOs involved in this pathway and summed across stations. Green bars indicate the fraction of transcript expression annotated to the genus whose absolute abundance profile is most highly correlated with the eigengene expression of the module concerned. Blue bars indicate the fraction of transcript expression annotated to the same taxonomic class as the genus represented by the green bars. Red bars indicate fraction of transcript expression annotated to other groups. Grey bars denote taxonomically unannotated transcripts.

**Supplementary Figure 11.**
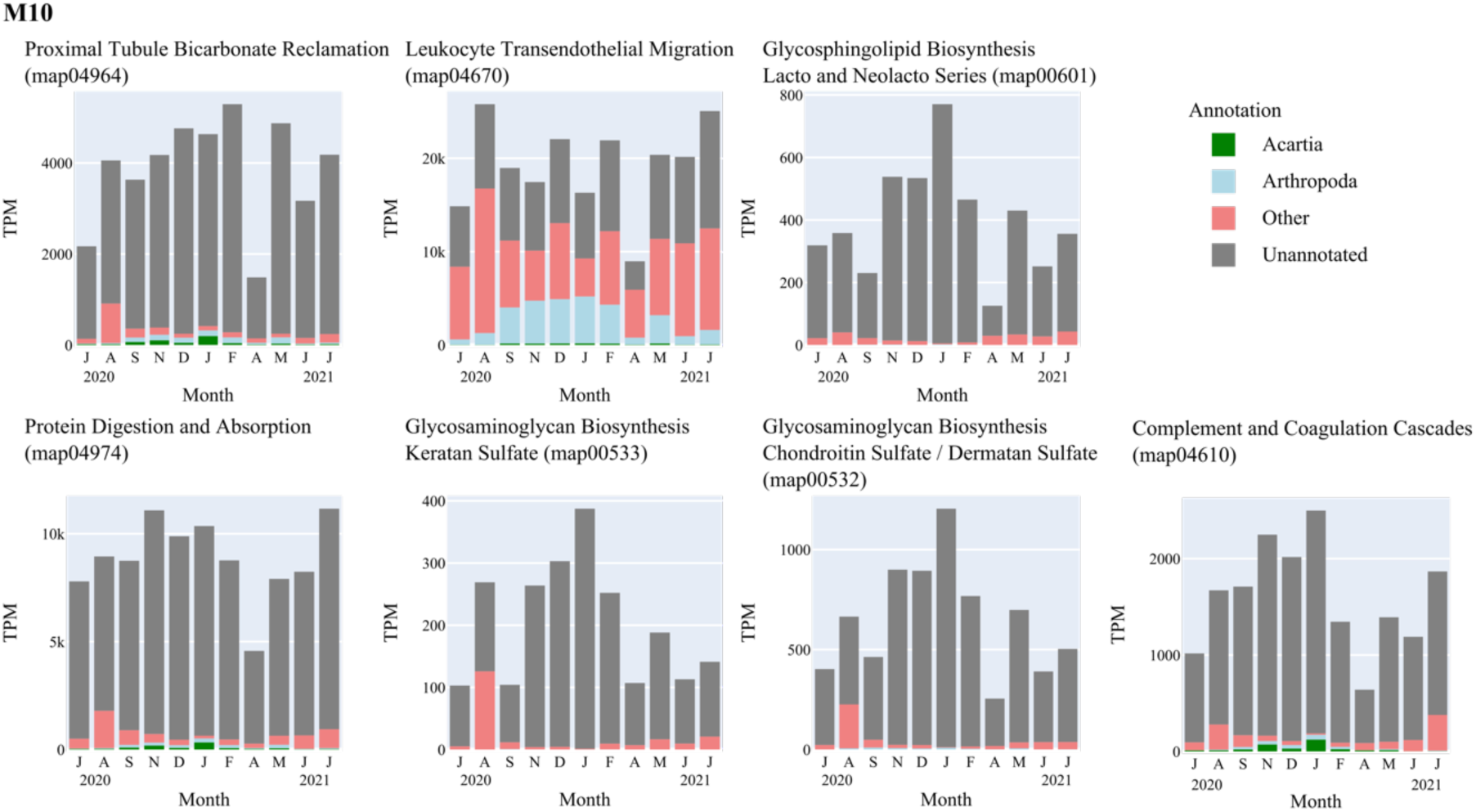
Expression of pathways characteristic of module 10. Characteristic pathways for module 10, as determined by Mann-Whitney U tests assessing the correlation between the TPM expression levels of a pathways’ KO identifiers and the eigengenes of the respective module. Each plot shows the monthly TPM values for a characteristic pathway, incorporating all transcripts annotated to KOs involved in this pathway and summed across stations. Green bars indicate the fraction of transcript expression annotated to the genus whose absolute abundance profile is most highly correlated with the eigengene expression of the module concerned. Blue bars indicate the fraction of transcript expression annotated to the same taxonomic class as the genus represented by the green bars. Red bars indicate fraction of transcript expression annotated to other groups. Grey bars denote taxonomically unannotated transcripts.

**Supplementary Figure 12.**
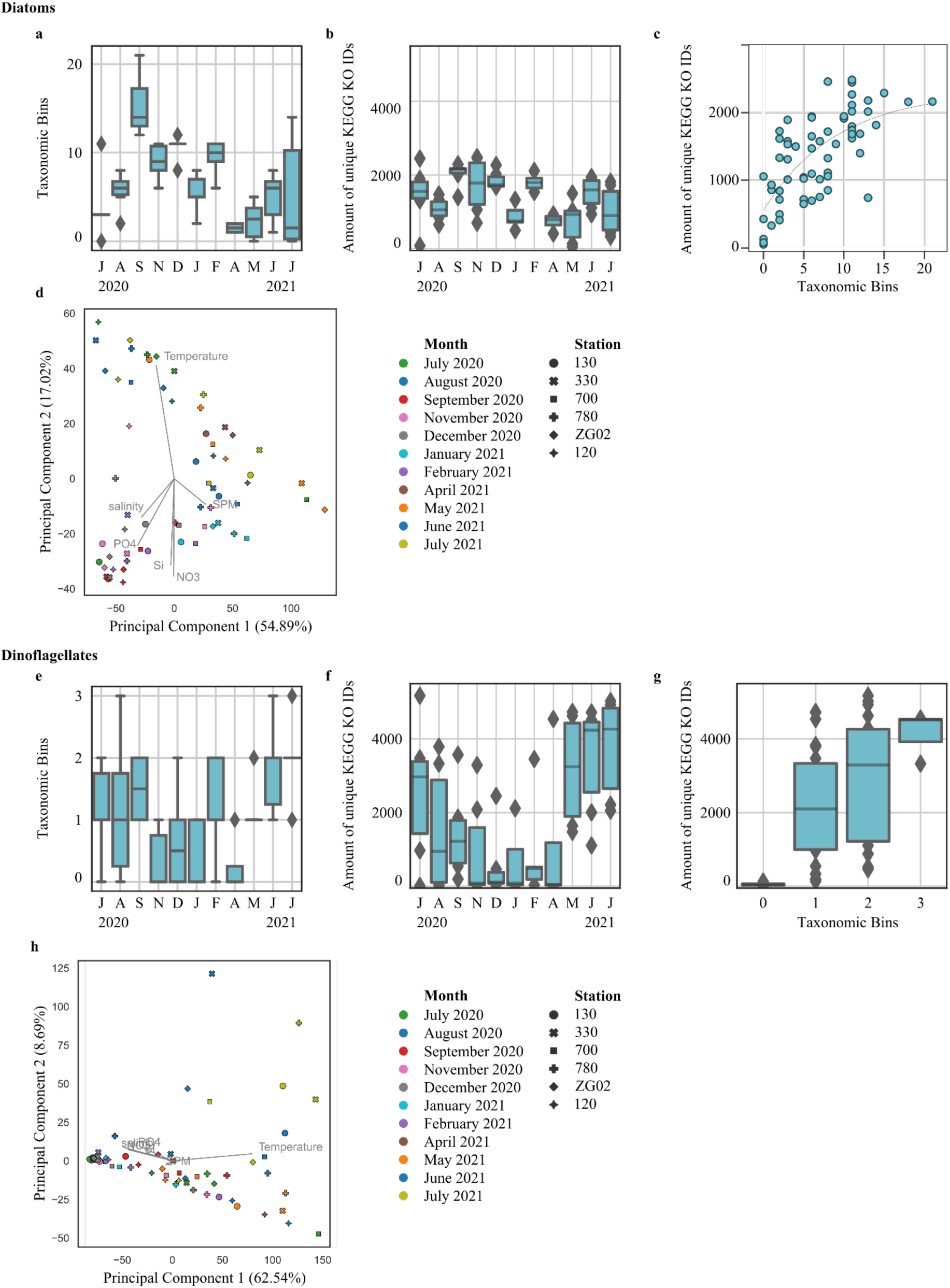
The occurrence of taxonomic bins, functional richness and functional diversity for diatoms and dinoflagellates annotated using EukProt. **a & e)** Boxplots of the number of active diatom and dinoflagellate species (taxonomic species bins with a minimum of 100 transcripts with non-zero expression in at least one sample) in a sample over time. Boxes encompass interquartile range, with the central line showing median. Diamonds represent outliers. **b & f)** Boxenplots of the number of unique KEGG KO IDs in a sample over time. Boxes encompass interquartile range, with the central line showing median. Diamonds represent outliers. **c)** Scatterplot depicting the relation between the amount of unique KEGG KO IDs and the number of active species. The grey line traces a fitted exponential model. **g)** Boxenplot of the number of unique KEGG KO IDs in relation to the number of active species. Boxes encompass interquartile range, with the central line showing median. Diamonds represent outliers. **d-h)** Principal Component Analyses on the log-transformed KEGG KO ID expression data from diatom and dinoflagellate EukProt annotated transcripts.

**Supplementary Figure 13.**
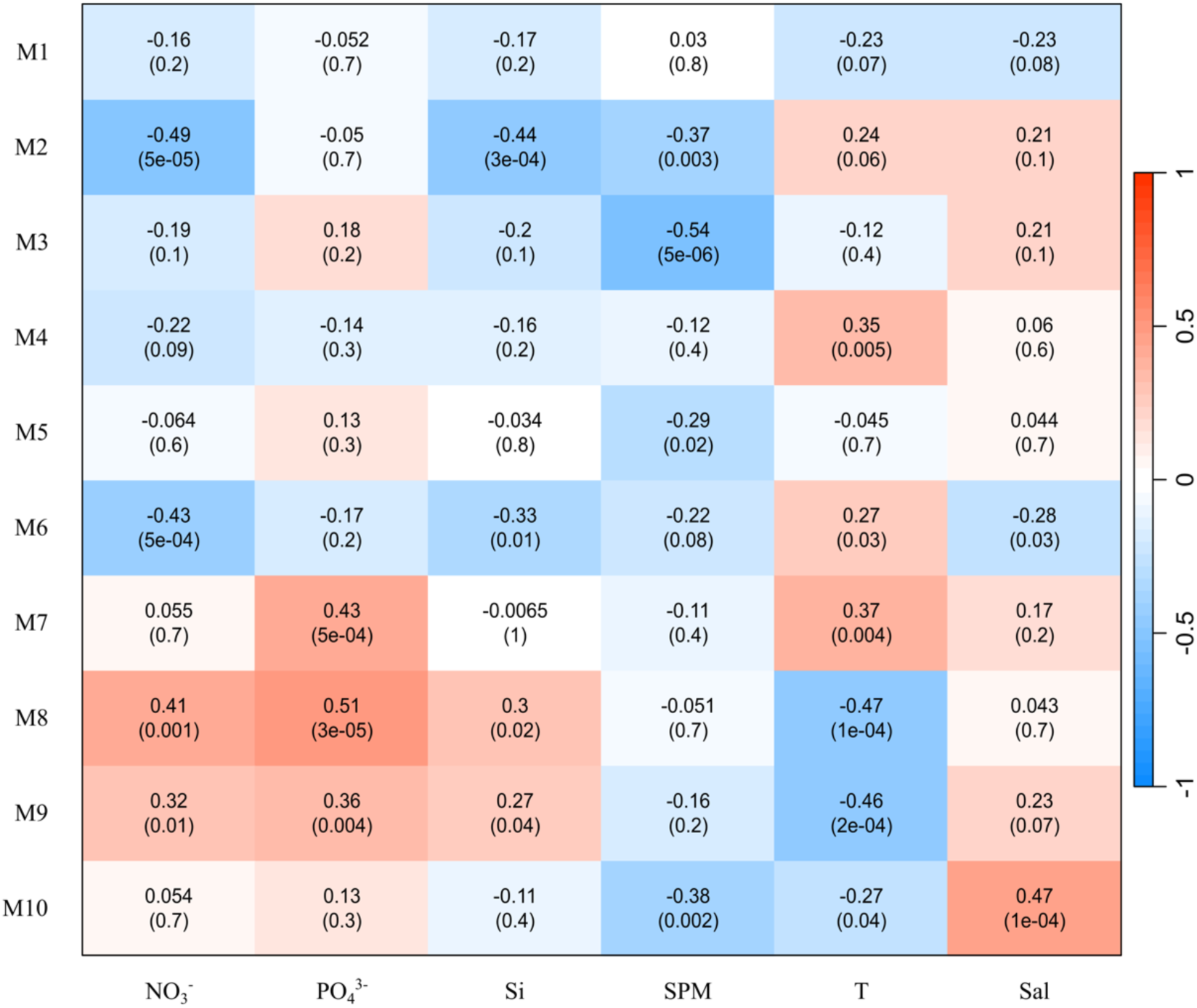
Correlation between module eigengene expression and environmental parameters. Numbers indicate Pearson’s correlations and their significance.

**Supplementary Figure 14.**
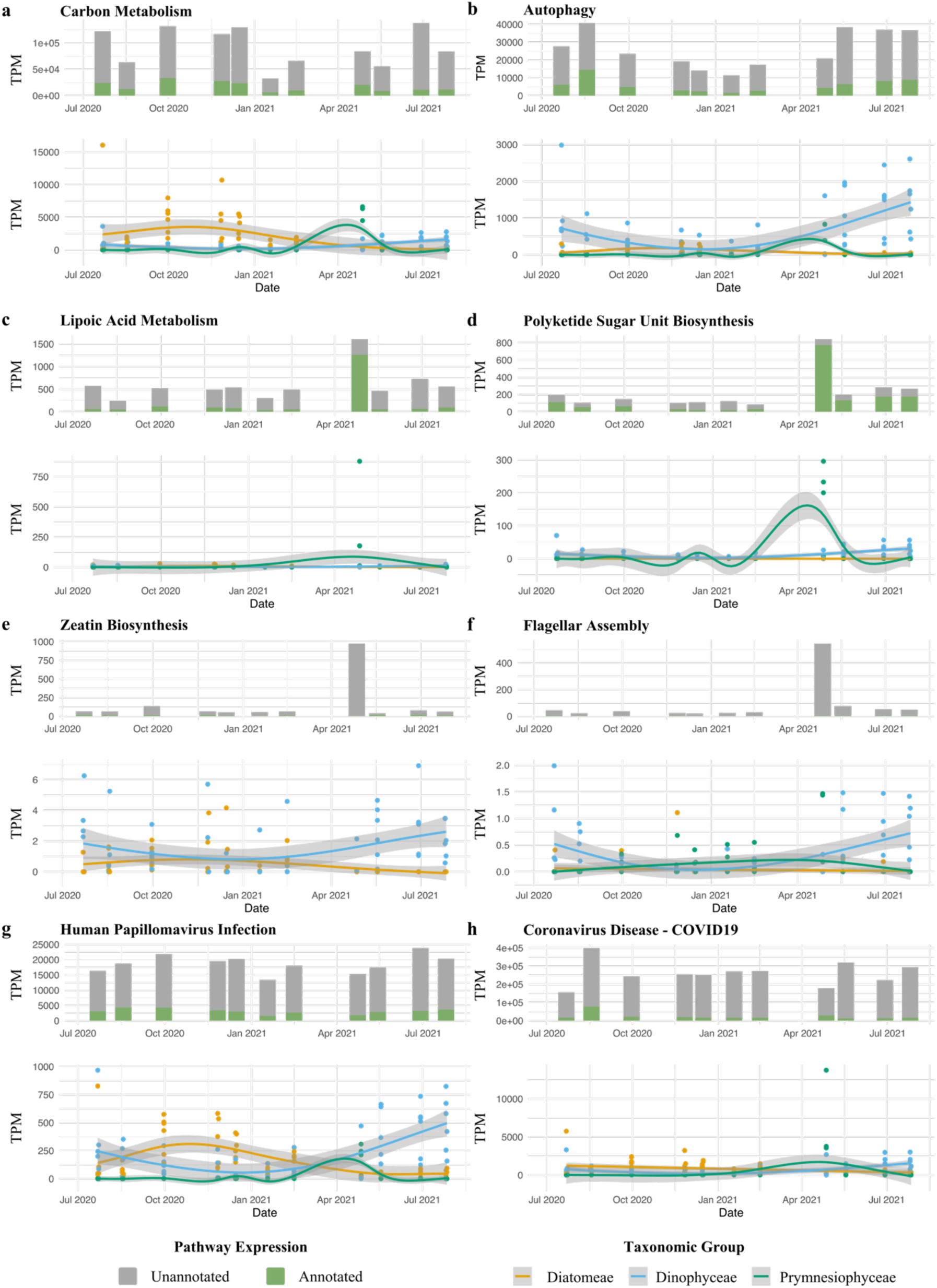
Expression of selected pathways and GAM regression fits for the expression of diatom, dinoflagellate, or *Phaeocystis* transcripts. Expression profiling of transcripts associated with a selection of KEGG pathways, alongside Generalised Additive Model (GAM) regression fits depicting the expression patterns of transcripts categorised within a respective pathway that were annotated as diatoms, dinoflagellates, or *Phaeocystis* (>90% sequence identity) over time. The top figure in a panel contrasts the expression levels of taxonomically annotated versus unannotated transcripts associated with the KEGG pathway. 8 pathways are displayed: **(a)** carbon metabolism, **(b)** autophagy, **(c)** lipoic acid metabolism, **(d)** polyketide sugar unit biosynthesis, **(e)** zeatin biosynthesis, **(f)** flagellar assembly, **(g)** human papillomavirus infection, and **(h)** coronavirus disease.

**Supplementary Figure 15.**
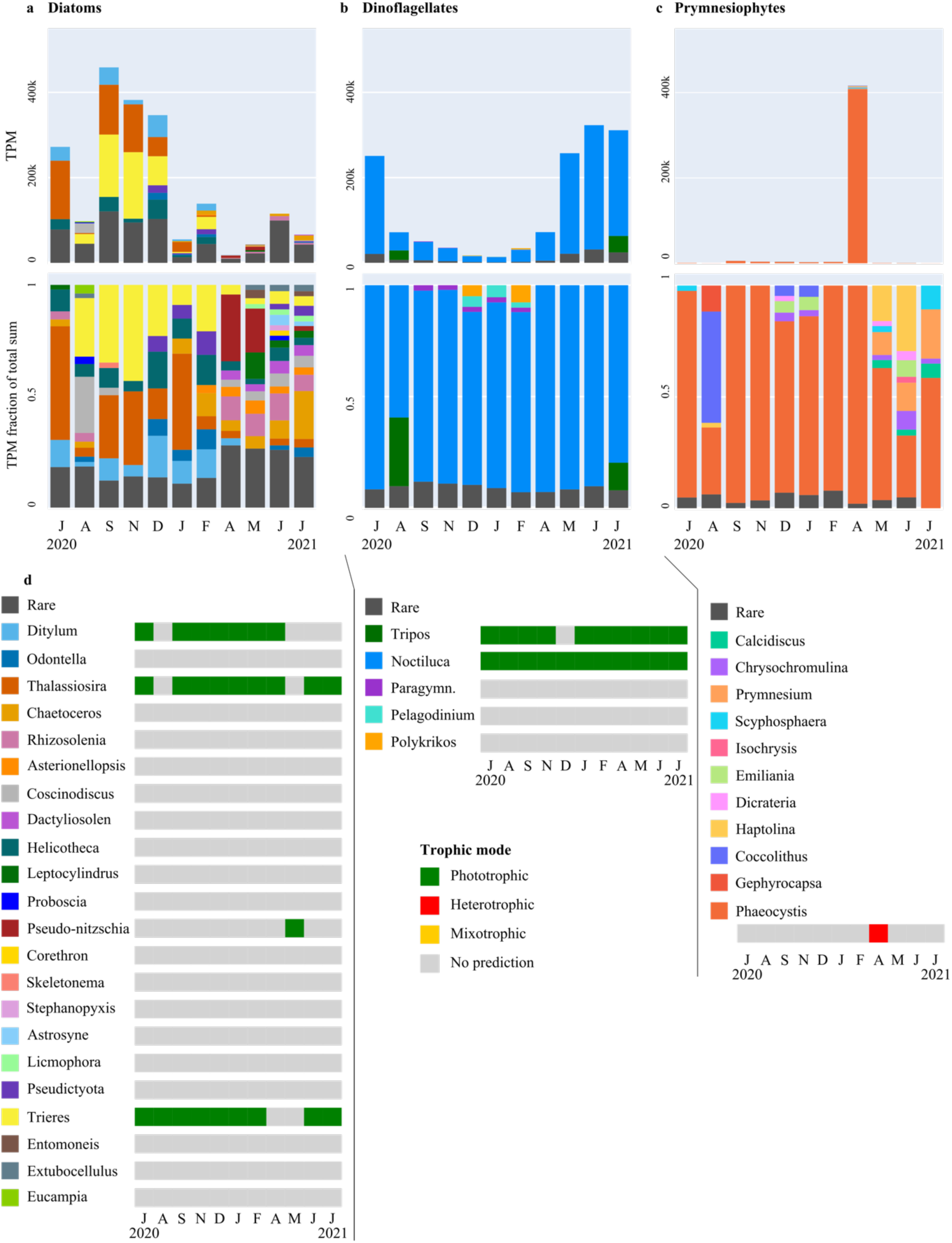
Absolute and relative turnover in diatom, dinoflagellate, and prymnesiophyte genera and their predicted trophic mode. **a)** The sum across sampling stations of transcript per million (TPM) counts attributed to diatom genera from July 2020 to July 2021 (top) and their relative taxonomic composition (bottom). **b)** The sum across sampling stations of TPM counts attributed to dinoflagellate genera from July 2020 to July 2021 (top) and their relative taxonomic composition (bottom). **c)** The sum across sampling stations of TPM counts attributed to prymnesiophyte genera from July 2020 to July 2021 (top) and their relative taxonomic composition (bottom). **d**) Trophic mode consensus prediction per species per month. Trophic modes could be predicted for taxonomic species bins that contained >800 PFAMs. The monthly consensus prediction was determined by a majority vote across samples (green: phototrophic, red: heterotrophic, grey: no prediction). Taxonomic annotations were obtained using EukProt.

**Supplementary Figure 16.**
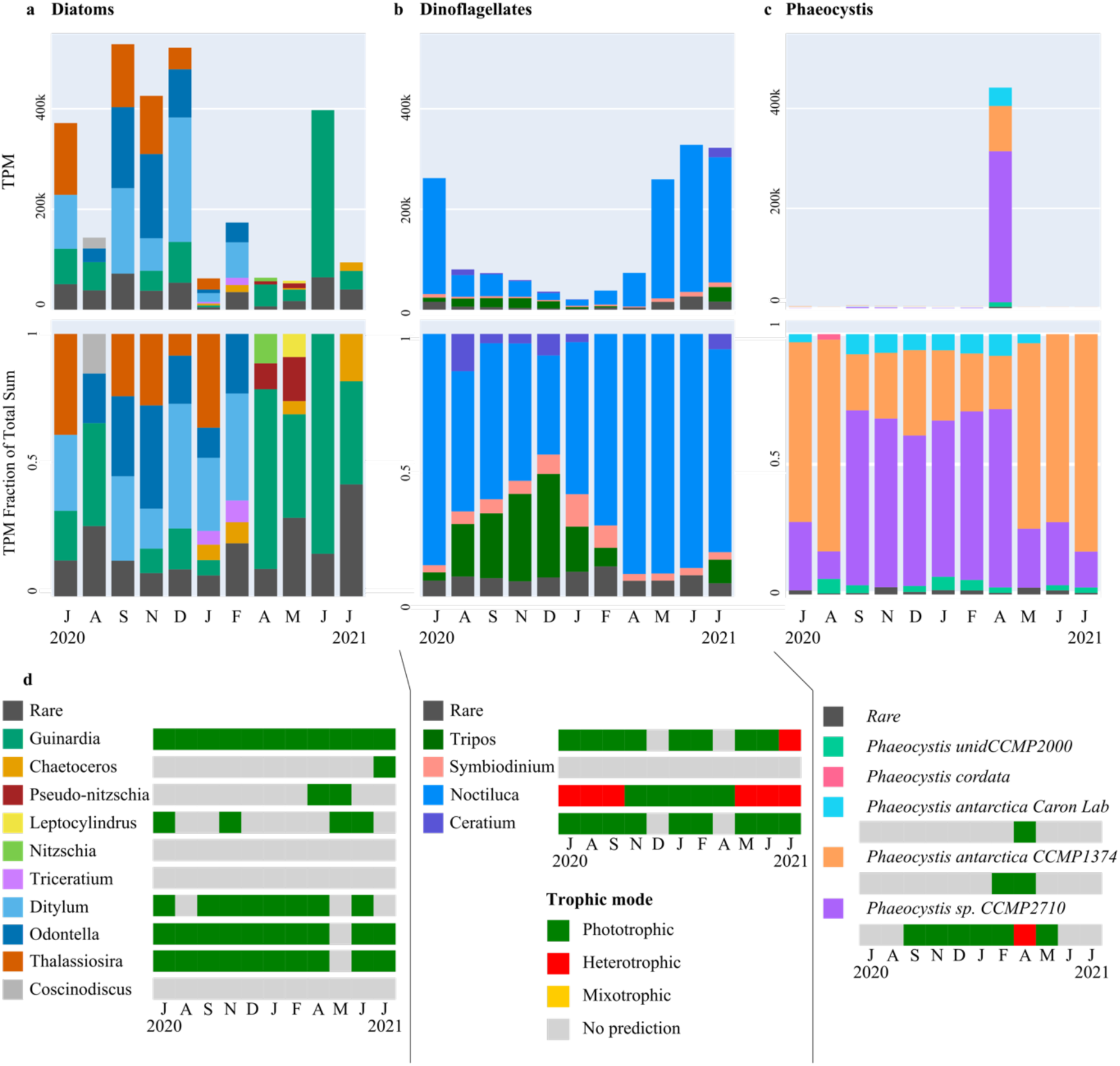
Absolute and relative turnover in diatom, dinoflagellate, and prymnesiophyte genera and their trophic mode based on PhyloDB taxonomic annotation. **a)** The sum of transcript per million (TPM) counts attributed to diatom genera from July 2020 to July 2021 and their relative taxonomic composition (bottom). **b)** The sum of TPM counts attributed to dinoflagellate genera from July 2020 to July 2021 and their relative taxonomic composition (bottom). **c)** The sum of TPM counts attributed to the genus *Phaeocystis* from July 2020 to July 2021 and their relative taxonomic composition (bottom). **d**) Trophic mode consensus prediction per month. For taxonomic species bins that contained >800 PFAMs, trophic mode could be predicted per sample. The monthly consensus prediction was determined by a majority vote across samples (green: phototrophic, red: heterotrophic, grey: no prediction). Taxonomic annotations were obtained using PhyloDB.

**Supplementary Figure 17.**
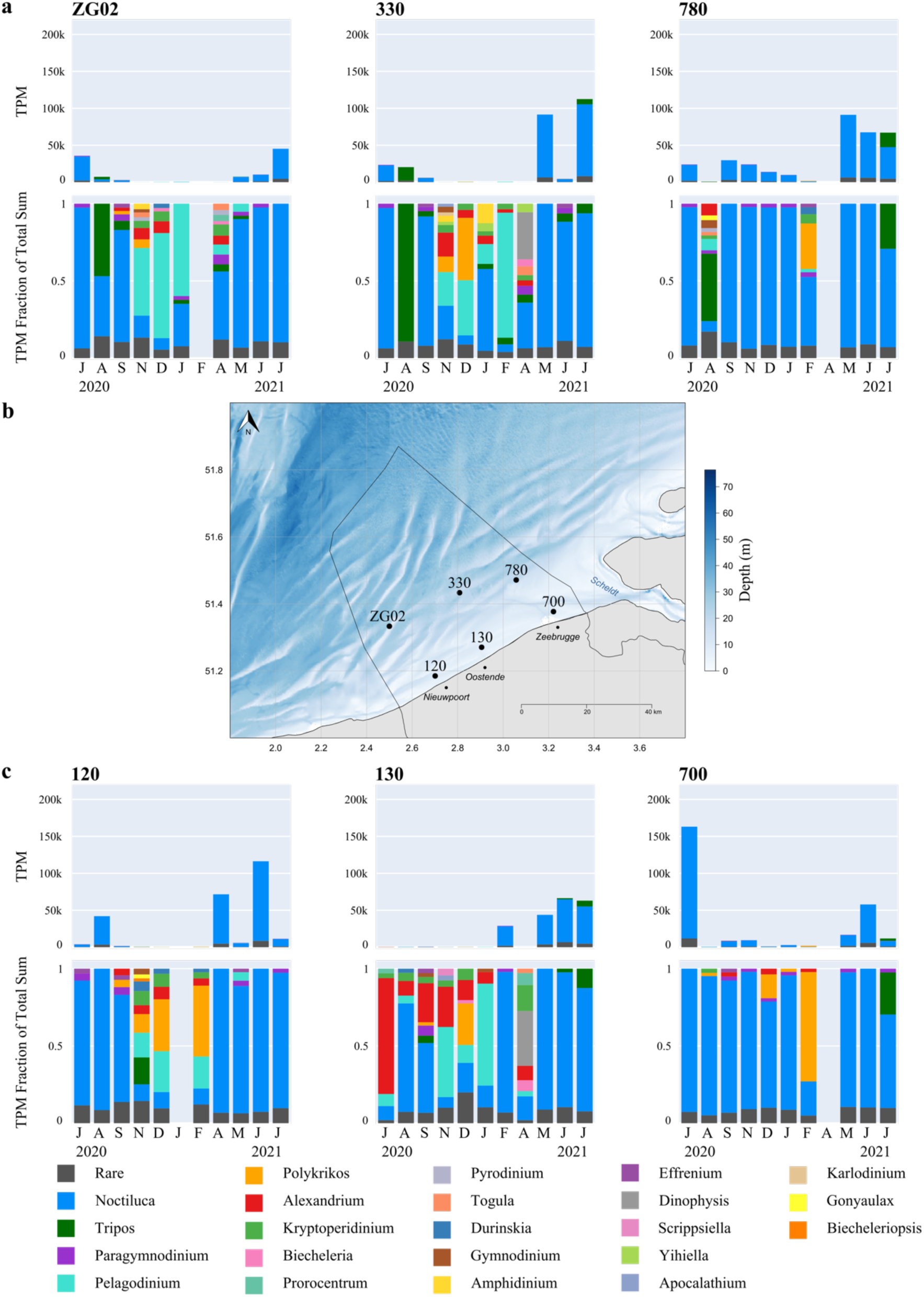
Monthly absolute and relative abundances of EukProt dinoflagellate genera at each station. **a)** Monthly transcript abundance and relative distribution of dinoflagellate genera annotated using EukProt, per offshore sampling station. Transcript abundance represents the sum of transcripts per million (TPM) belonging to a taxonomic group. Relative abundance was calculated as the sum of TPM for that group for a given sample, divided by the total TPM of all genera found in that sample. When relative abundance of a genus was <2 %, they were labelled as ‘rare’. **b)** Spatial location of the 6 sampling stations in the Belgian Part of the North Sea. c) Monthly transcript abundance and relative distribution of dinoflagellate genera annotated using EukProt, per nearshore sampling station. Transcript abundance and relative abundance were calculated the same as for the offshore stations.

**Supplementary Figure 18.**
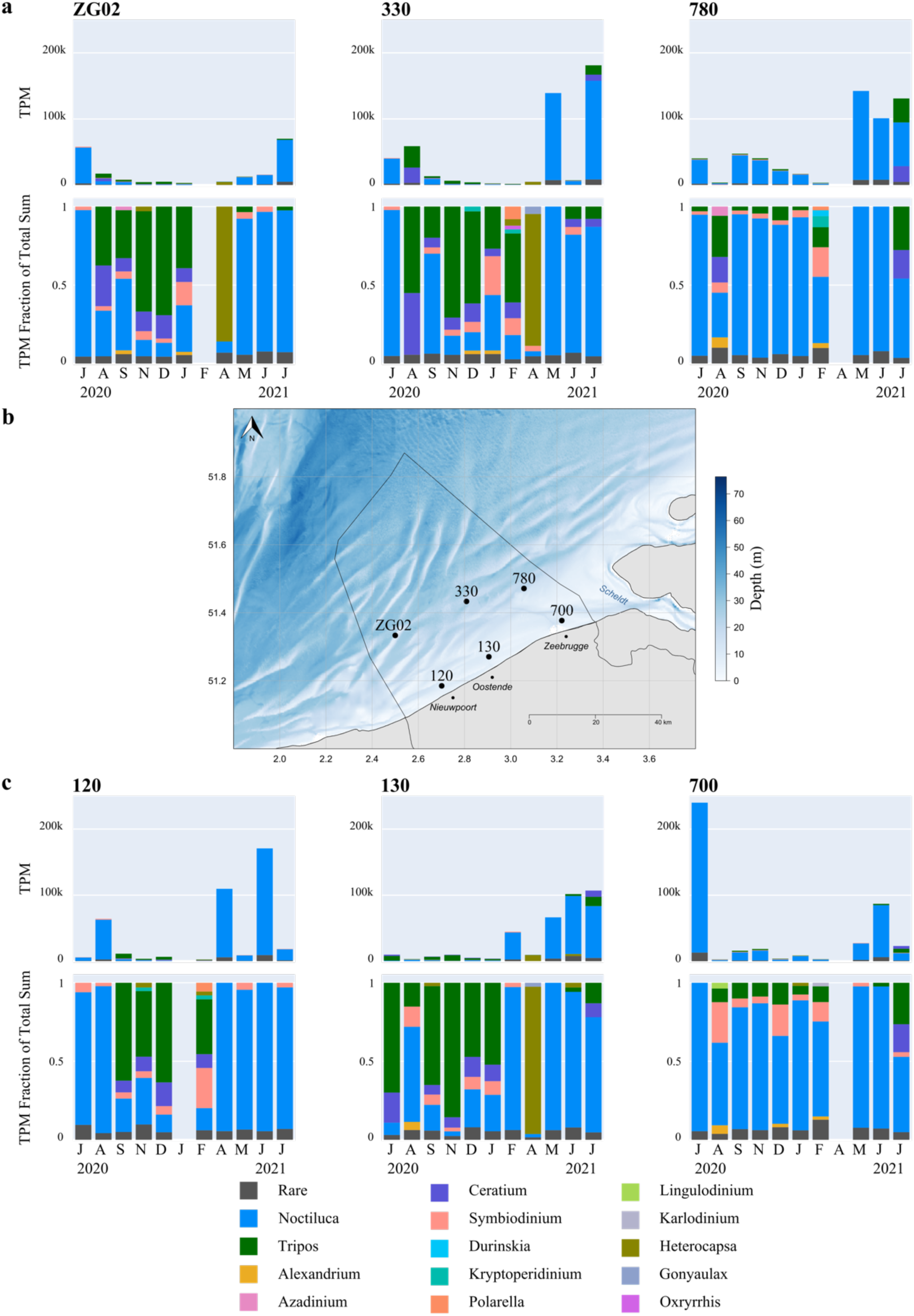
Monthly absolute and relative abundances of PhyloDB dinoflagellate genera at each station. **a)** Monthly transcript abundance and relative distribution of dinoflagellate genera annotated using PhyloDB, per offshore sampling station. Transcript abundance represents the sum of transcripts per million (TPM) belonging to a taxonomic group. Relative abundance was calculated as the sum of TPM for that group for a given sample, divided by the total TPM of all genera found in that sample. When relative abundance of a genus was <2 %, they were labelled as ‘rare’. **b)** Spatial location of the 6 sampling stations in the Belgian Part of the North Sea. c) Monthly transcript abundance and relative distribution of dinoflagellate genera annotated using PhyloDB, per nearshore sampling station. Transcript abundance and relative abundance were calculated the same as for the offshore stations.

**Supplementary Figure 19.**
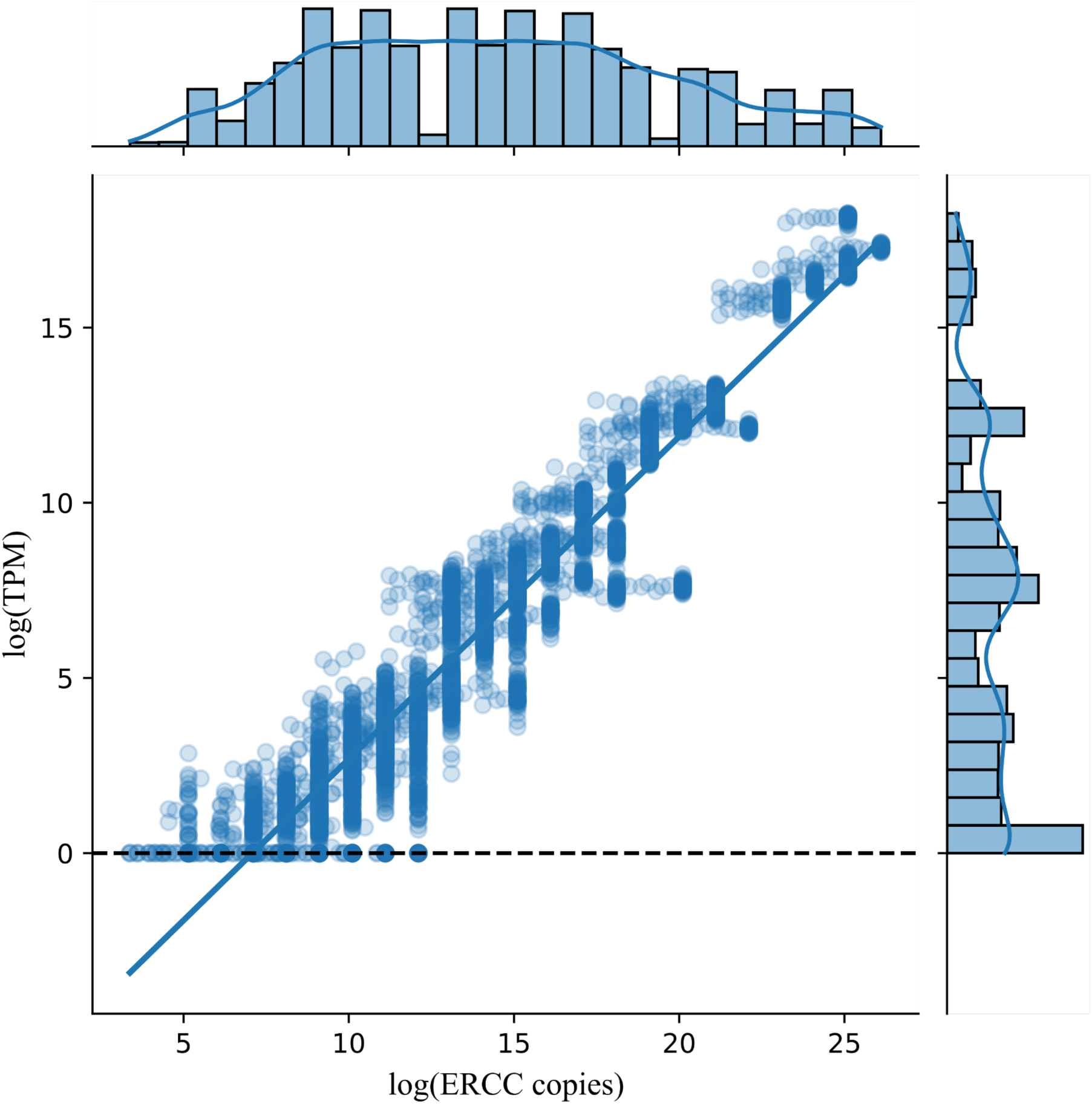
The relationship between ERCC92 standards and their associated TPM counts. Jointplot illustrating the relationship between log2-transformed ERC992 transcript standards copies and their corresponding log-transformed transcript per million counts. The fitted trendline helps determine the lower limit of detection of expression quantification.

### Supplementary Tables

**Supplementary Table 1.**
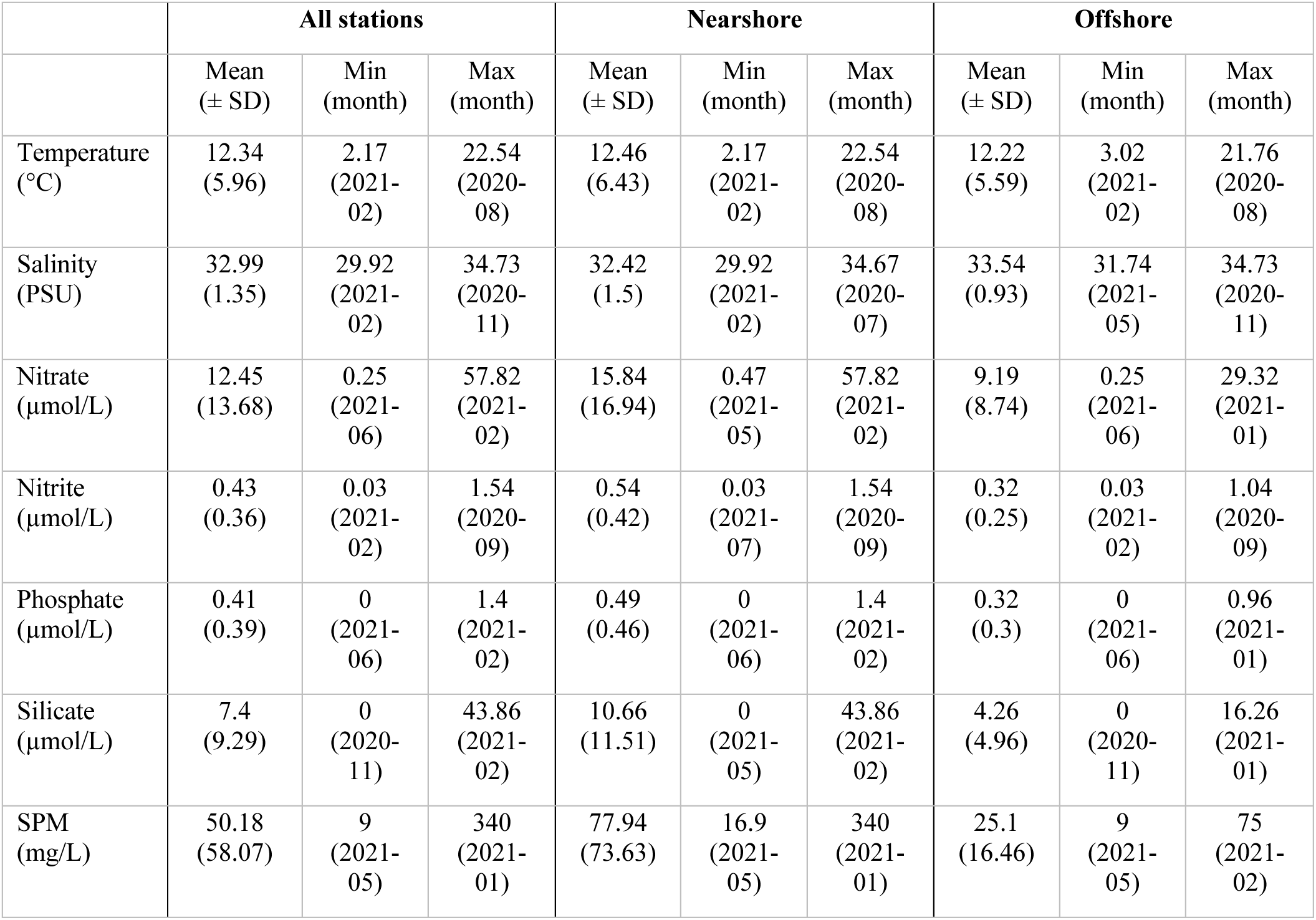
Summary of abiotic environmental variables measured in the Belgian Part of the North Sea. This table presents mean values of temperature, salinity, nitrate, nitrite, phosphate, silicate, and suspended particulate matter (SPM) data for all, nearshore (station 120, 130, and 700), and offshore (station ZG02, 330, and 780) stations.

|  | All stations |  |  | Nearshore |  |  | Offshore |  |  |
| --- | --- | --- | --- | --- | --- | --- | --- | --- | --- |
| | Mean<br>(± SD) | Min<br>(month) | Max<br>(month) | Mean<br>(± SD) | Min<br>(month) | Max<br>(month) | Mean<br>(± SD) | Min<br>(month) | Max<br>(month) |
| Temperature<br>(°C) | 12.34<br>(5.96) | 2.17<br>(2021-02) | 22.54<br>(2020-08) | 12.46<br>(6.43) | 2.17<br>(2021-02) | 22.54<br>(2020-08) | 12.22<br>(5.59) | 3.02<br>(2021-02) | 21.76<br>(2020-08) |
| Salinity<br>(PSU) | 32.99<br>(1.35) | 29.92<br>(2021-02) | 34.73<br>(2020-11) | 32.42<br>(1.5) | 29.92<br>(2021-02) | 34.67<br>(2020-07) | 33.54<br>(0.93) | 31.74<br>(2021-05) | 34.73<br>(2020-11) |
| Nitrate<br>(μmol/L) | 12.45<br>(13.68) | 0.25<br>(2021-06) | 57.82<br>(2021-02) | 15.84<br>(16.94) | 0.47<br>(2021-05) | 57.82<br>(2021-02) | 9.19<br>(8.74) | 0.25<br>(2021-06) | 29.32<br>(2021-01) |
| Nitrite<br>(μmol/L) | 0.43<br>(0.36) | 0.03<br>(2021-02) | 1.54<br>(2020-09) | 0.54<br>(0.42) | 0.03<br>(2021-07) | 1.54<br>(2020-09) | 0.32<br>(0.25) | 0.03<br>(2021-02) | 1.04<br>(2020-09) |
| Phosphate<br>(μmol/L) | 0.41<br>(0.39) | 0<br>(2021-06) | 1.4<br>(2021-02) | 0.49<br>(0.46) | 0<br>(2021-06) | 1.4<br>(2021-02) | 0.32<br>(0.3) | 0<br>(2021-06) | 0.96<br>(2021-01) |
| Silicate<br>(μmol/L) | 7.4<br>(9.29) | 0<br>(2020-11) | 43.86<br>(2021-02) | 10.66<br>(11.51) | 0<br>(2021-05) | 43.86<br>(2021-02) | 4.26<br>(4.96) | 0<br>(2020-11) | 16.26<br>(2021-01) |
| SPM<br>(mg/L) | 50.18<br>(58.07) | 9<br>(2021-05) | 340<br>(2021-01) | 77.94<br>(73.63) | 16.9<br>(2021-05) | 340<br>(2021-01) | 25.1<br>(16.46) | 9<br>(2021-05) | 75<br>(2021-02) |

## The North Sea micro-eukaryotic metatranscriptome

A total of 1.049 billion raw reads were generated from sea surface water samples, an average of 16 million (SD=2.3M) per sample. The resulting de novo metatranscriptome assembly contained over 7 million unique transcripts with a median transcript length of 342 bp (Supplementary Fig. 1). 3,705,883 proteins were predicted from the assembled transcripts. Functional or taxonomic annotation information was found for 79% of predicted proteins. 2,235,576 proteins, 60% of the total, could be functionally annotated (Supplementary Fig. 1c & d). Shallow taxonomic annotations, here defined as 60% sequence identity with either the PhyloDB or the EukProt reference databases, were found for 59% and 64% of proteins respectively. When using a stricter cut-off value of 90% sequence identity to obtain deeper taxonomic resolution, 26% of proteins could be assigned a taxonomic identity using PhyloDB and 13% matched with a EukProt reference sequence. Given the broader representation of eukaryotic diversity and more recent release date, we used the EukProt reference database for further analyses. However, for specific taxonomic groups, such as diatoms and dinoflagellates, both databases were consulted to obtain a more comprehensive picture. To further assess the representation of southern North Sea species in other global ocean reference databases, we examined the alignment of our assembled transcripts to the Tara Oceans’ metagenome assembled genomes (MAGs) database^1^ (retaining only alignments with at least 80% coverage and 95% sequence identity). On average, 2.59% of the assembled transcripts mapped to the collection of Tara oceans’ eukaryotic genomes. For April samples, however, we observed higher mapping rates with 14% of the assembled transcripts mapping to the MAGs, due to high mapping rates against *Phaeocystis* genomes. The overall low taxonomic representation in the MAGs database indicates an underrepresentation of microeukaryotic plankton species from our study area in global ocean reference databases. This highlights the importance of sampling, extracting, and characterising genomic data from the southern North Sea and similar temperate coastal marine ecosystems.

## Supplementary Methods

### Calculating cell densities from FlowCam image data

To obtain microphytoplankton biomass estimates, 50 L of sea surface water was collected using a stainless-steel bucket and filtered through a 55 µm mesh size Apstein net. Filtered samples were transferred to a plastic falcon and fixed with acidic Lugol’s iodine solution (1-5 %). Samples were then stored in the dark at 4 °C until laboratory processing. In the lab, samples were processed using the FlowCam VS-4 bench-top model (Fluid Imaging Technologies, Yarmouth, Maine, U.S.A.) equipped with a Sony XCD SC90 digital grey-scale camera and VisualSpreadsheet® software (version 4.2.52). Samples were prefiltered at 300µm to avoid clogging of flowcells and a predefined threshold ensured only particles in the 50-300µm range were captured. A first presample run was used to determine particle load and determine sample dilution factors if Particles Per Used Image exceeded values of 1.2. Three replicate runs of each sample were carried out to minimise technical variation. The resulting output, consisting of collages of particle images, a set of image parameters, and sampling and processing metadata, were processed and uploaded to the internal BioSens MongoDB database. Unvalidated single images were then fed to a convolutional neural network, trained on an extensive dataset of human-validated images, that classified each particle into one of over 90 distinct taxonomic groups^2^. All model predictions were verified by human taxonomists.

Cell densities of the assigned taxonomic groups were calculated using the filtered volume of sea surface water, the volume of the sample measured in the lab, the fluid volume imaged by FlowCam, the groups’ particle count, and a dilution factor calculated from the dilution of samples:

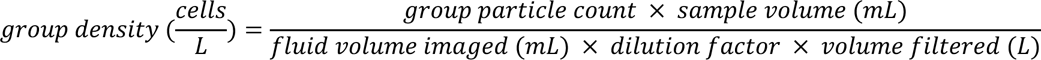

The resulting cell densities were checked for possible outliers. The validated FlowCam data is made available through the Belgian LifeWatch RShiny application at https://rshiny.vsc.lifewatch.be/flowcam-data/. For more detailed information on the full FlowCam protocol see Martínez et al. (2020)^2^.

### Transcript and ERCC spike quantification & normalisation

Transcript and ERCC spike-in abundances were quantified using Kallisto, yielding TPM (Transcript Per Million) counts for every transcript^3^. Using the known molar concentrations of the 92 RNA spikes, a lower limit of detection (LLD) could be calculated (Fig. S19). The LLD is the concentration of a given transcript (attomoles/µL), or the number of copies, that needs to be present in a sample to yield one TPM count. TPM counts below the trusted LLD were set to We calculated transcripts per L using ERCC RNA spike-in recovery values according to:

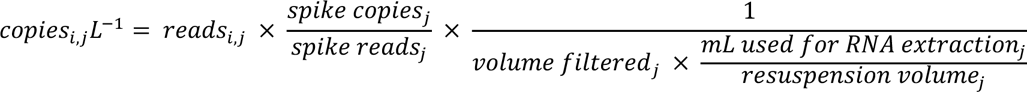

Where *Copies_i,j_L^-1^* equals the amount of RNA copies per L of seawater of transcript *i* in sample *j*, *reads_i,j_* are the TPM values of read *i* in sample *j*, spike copies*_j_* is the average amount of copies added to sample *j*, spike reads*_j_* are the average amount of spike reads (TPM) in sample *j*, volume filtered*_j_* is the volume of seawater filtered for sample *j*, and resuspension volume*_j_* is the amount of seawater used to re-elute the filtration residue^4^. In calculating the amount of RNA copies per L of seawater we ignored the loss of extracted RNA for quality control, as these volumes were equal for all samples.

### WGCNA

A weighted gene co-expression network analysis (WGCNA) was performed using the WGCNA package^5^ (version 1.72-1) in R^6^. The approach was modified from the analysis pipeline described in Cohen et al, 2021^7^. Briefly, we aimed to identify co-expression modules of Kyoto Encyclopedia of Genes and Genomes (KEGG) KO identifiers from log-normalised count sums of KEGG KOs over annotated transcripts. The expression matrix was pre-processed by setting TPM values below 1 to 0 and removing identifiers with total counts below 10 across samples. Outlier samples were detected using the WGCNA scaled connectivity measure, and the sample dendrograms were constructed using average linkage hierarchical clustering on the Pearson correlation (PC) network of KEGG KOs. Module detection was performed using the WGCNA dynamic tree cut algorithm (minimum module size: 70, deepSplit = 4), and modules with highly similar expression profiles (PC>0.6) were merged. Module eigengenes, representing the first principal component of each module, were calculated using the WGCNA package moduleEigengene function (version 1.72-1), and their correlations with environmental parameters and the absolute TPM abundances of taxonomic groups were assessed. To identify metabolic pathways associated with a given module eigengene, we performed Mann-Whitney U tests using SciPy’s mannwhitneyu function to assess which pathways were more represented at the top of a ranked list of KEGG KO identifiers, ordered according to decreasing module membership, than expected by chance^8^. Module membership of a pathway was defined as the Pearson correlation between module eigengene expression and KEGG KO identifier. Resulting p-values were adjusted for multiple testing with the Benjamini-Hochberg procedure, which controls the false discovery rate^9^.

## Notes

https://dataview.ncbi.nlm.nih.gov/object/PRJNA1021244

https://github.com/MichielPerneel/BPNS_seasonal_MTX

### Competing Interest Statement

The authors have declared no competing interest.

### Summary of Updates

Fixed error in author names.

